# Maternal SETDB1 enables development beyond cleavage stages by extinguishing the MERVL-driven 2-cell totipotency transcriptional program in the mouse embryo

**DOI:** 10.1101/2025.10.07.680892

**Authors:** Tie-Bo Zeng, Zhen Fu, Mary F. Majewski, Ji Liao, Marie Adams, Piroska E. Szabó

## Abstract

Loss of maternal SETDB1, a histone H3K9 methyltransferase, leads to developmental arrest prior to implantation, with very few mouse embryos advancing beyond the 8-cell stage, which is currently unexplained. We genetically investigate SETDB1’s role in the epigenetic control of the transition from totipotency to pluripotency—a process demanding precise timing and forward directionality. Through single-embryo total RNA sequencing of 2-cell and 8-cell embryos, we find that *Setdb1*^mat-/+^ embryos fail to extinguish 1-cell and 2-cell transient genes—alongside persistent expression of MERVL retroelements and MERVL-driven chimeric transcripts that define the totipotent state in mouse 2-cell embryos. Comparative bioinformatics reveals that SETDB1 acts at MT2 LTRs and MERVL-driven chimeric transcripts, which normally acquire H3K9me3 during early development. The dysregulated targets substantially overlap with DUXBL-responsive genes, indicating a shared regulatory pathway for silencing the 2-cell transcriptional program. We establish maternal SETDB1 as a critical chromatin regulator required to extinguish retroelement-driven totipotency networks and ensure successful preimplantation development.

## Introduction

SETDB1 is a histone 3 lysine 9 (H3K9) di- and tri-methyl transferase [1]. In the mouse embryo, it localizes to the inner cell mass of the blastocyst [2] and is essential for postimplantation development [3]. Homozygous knockout of *Setdb1* results in embryonic lethality between 3.5- and 5.5-days post coitum (dpc) and prevents the derivation of embryonic stem cells (ESCs). SETDB1 is required for the silencing of developmental gene promoters [4]. Although *Setdb1*^-/-^ zygotic mutants retain maternally deposited SETDB1 protein from the oocyte, conditional knockout of *Setdb1* during oogenesis produces an earlier and more severe maternal-effect phenotype: most oocytes arrest during meiosis, and the few that are fertilized typically fail to develop beyond the 2-cell or 8-cell stages and never reach the blastocyst [5, 6]. The basis of this cleavage-stage arrest in the *Setdb1*^mat-/+^ embryos remains unknown.

This developmental block coincides with the transition from totipotency to pluripotency, prompting us to investigate whether SETDB1 regulates this critical process. Following fertilization, parental epigenomes undergo extensive reprogramming to enable zygotic genome activation (ZGA). In mice, minor ZGA occurs at the late 1-cell stage and major ZGA at the 2-cell stage—both essential for development [7]. The 2-cell stage marks the onset of totipotency, while from the 4-cell stage onward, blastomeres begin committing to either the pluripotent inner cell mass or the trophectoderm lineage [8]. In ESCs, dedifferentiation to 2-cell-like cells (2CLCs) [9] depends on DUX, a pioneer transcription factor that activates ZGA genes [10]. Additional ZGA drivers include OBOX, NFYA, NR5A2, and SRF [10–15], which open chromatin and activate transcription in a combinatorial or compensatory fashion [16]. Although DUX and individual OBOX proteins are dispensable for ZGA in embryos [11, 14, 17, 18], their combined loss [14], or loss of both DUX and OBOX4 [19], impairs development, suggesting functional redundancy. While recent studies shed light on ZGA initiation, the mechanisms responsible for extinguishing the 2-cell transcriptional program remain poorly understood.

Endogenous retroviruses (ERVs) are key regulators of early development. MERVL elements are transcriptionally active in 2-cell embryos [20] and drive chimeric transcripts that define the totipotent state [9, 21]. MERVL activation is required for blastocyst formation [22] and is mediated by DUX/OBOX and SRF binding at MT2 long terminal repeats (LTRs) [13, 23]. However, sustained MERVL expression—such as that induced by forced DUX expression—disrupts totipotency exit and embryo development [18, 24]. Although timely repression of MERVLs and the 2-cell program requires DUXBL [25], the mechanisms by which DUXBL silences totipotency-related genes remain unknown.

Because genome-wide DNA demethylation occurs after fertilization, preimplantation embryos rely heavily on histone modifications to regulate transcription. During cleavage-stage development, H3K9me3 is deposited at retrotransposons; MERVL elements begin to acquire this repressive mark by the 2-cell stage, as DNA methylation levels decline [26]. SETDB1, a key H3K9 methyltransferase, is thus a strong candidate for regulating MERVL silencing and early embryonic transcriptional control. In ESCs, SETDB1 represses transposable elements (TEs) via TRIM28 (KAP1)-dependent H3K9me3 deposition and by maintaining DNA methylation at LTRs and imprinted DMRs [27–29]. However, its role at MERVLs in ESCs is debated: some studies report minimal repression [30], while others show that SETDB1 silences MERVLs to prevent dedifferentiation into 2CLCs [31]. In oocytes, *Setdb1* loss leads to derepression of ERVK and ERVL-MaLR retrotransposons [5, 6]. Whether SETDB1 also regulates MERVLs and totipotency exit in cleavage-stage embryos remains unresolved.

To address this question and understand why *Setdb1*^mat-/+^ embryos fail to exit the cleavage stages, we employed a conditional knockout strategy. We previously showed that maternal SETDB1 protects the maternal pronucleus from TET3-mediated DNA demethylation in zygotes [32]. Here, we investigate its role in preimplantation development by analyzing the transcriptomes of 2-cell and rare 8-cell *Setdb1*^mat-/+^ embryos. We find that maternally deposited SETDB1 is required to silence MERVL expression and 2-cell-specific totipotency transcripts, enabling successful progression beyond the cleavage stages.

## Results

### *Setdb1* maternal knockout embryos fail preimplantation development

A complete understanding of early embryonic development—and its intrinsic directionality and tempo—requires in vivo approaches. To address the in vivo role of SETDB1 in controlling the earliest stages of embryo development we applied a genetic approach. We crossed *Setdb1*^f/f^; *Zp3*-cre females with wild-type JF1/Ms males to obtain maternal knockout (*Setdb1*^mat-/+^) (KO) embryos. Control *Setdb1*^f/f^ (WT) embryos were derived from *Setdb1*^f/f^ females lacking the *Zp3*-cre transgene. We previously reported that H3K9me2 and H3K9me3 histone marks are globally reduced in *Setdb1*^mat-/+^ zygotes following *Zp3*-cre -driven excision of the SET domain from growing oocytes [32].

The maternal *Setdb1* transcript is present in oocytes and persists throughout preimplantation development, while zygotic expression from the paternal allele begins only at the blastocyst stage [3]. Consequently, *Setdb1*^mat-/+^ embryos lack SETDB1 protein function from the oocyte through the 8-cell stage.

At 1.5 dpc, over 70% of control eggs had reached the 2-cell (2c) stage, with a few progressing to the 4-cell (4c) stage (Fig. 1A). In contrast, fewer than 30% of *Setdb1*^mat-/+^ eggs reached the 2c stage, with many arrested at GV, MI, or MII stages. By 2.5 dpc, ∼80% of control embryos reached the 8-cell (8c) stage, while only ∼4% of mutant embryos did so. These findings agree with earlier studies [5, 6] and show that preimplantation development is severely impacted in *Setdb1*^mat-/+^ embryos.

**Figure 1.**
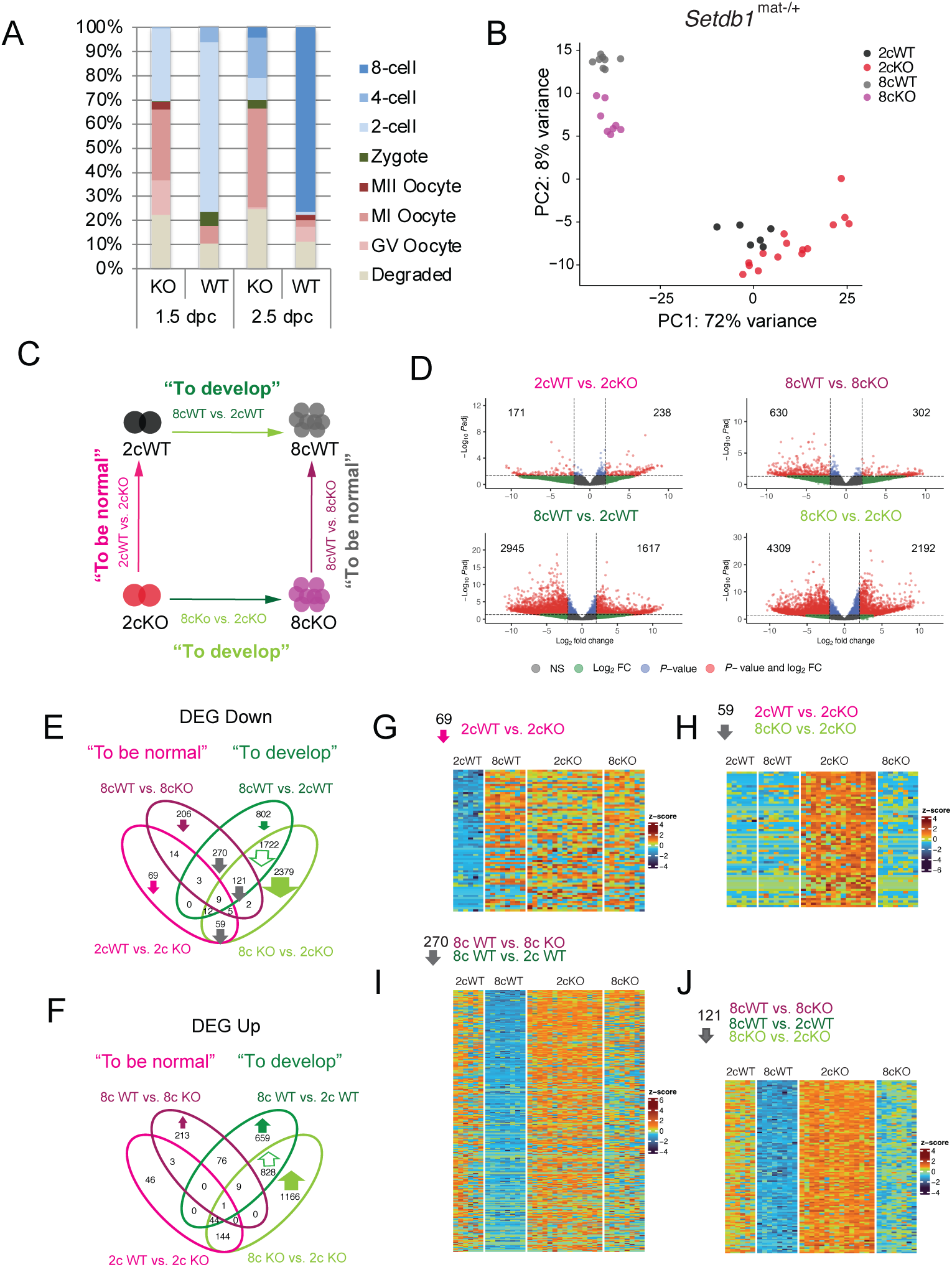
Maternal SETDB1 is essential for development beyond the 8-cell stage. (A) Quantification of *Setdb1*^mat-/+^ (KO) and *Setdb1*^fl/+^ (WT) embryo stages from the following number of total recovered embryos: KO (n=638), WT (n=484) at 1.5 dpc, and KO (n=310) and WT (n=80) at 2.5 dpc. (B) Principal component analysis of single-embryo total RNA-seq data from 2cWT (n=6), 2cKO (n=15), 8cWT (n=8), and 8cKO (n=8) embryos. (C) Schematic of four pair-wise comparisons defining requirements for normalcy and development. (D) Volcano plots highlighting DEGs using |log₂FC| > 1 and adjusted *P* < 0.05. (E–F) Four-way DEG comparisons visualized by Venn diagrams: (E) downregulated; (F) upregulated. (G–J) Heatmaps of DEGs from Venn compartments, showing stage- and genotype-specific patterns.

### Single-embryo total RNA sequencing of *Setdb1*^mat-/+^ embryos

As no *Setdb1*^mat-/+^ embryos develop into blastocysts, we performed single-embryo RNA sequencing at the 2c and 8c stages (1.5 and 2.5 dpc, respectively) to determine how maternal SETDB1 supports development beyond the cleavage stage. The KO embryos that did develop to the 2c or 8c stage were morphologically indistinguishable from WT under light microscopy (Figure S1A). To optimally assess both regular and chimeric transcription we used total RNA sequencing. We profiled both KO and WT embryos with >6 biological replicates (Table S1). Examples of normally expressing transcripts are displayed in Figure S1B.

Principal component analysis (PCA) revealed that samples clustered primarily by developmental stage (PC1), and secondarily by genotype (PC2) (Figure 1B), indicating that stage-specific transcriptional programs remain partially impact—more distinctly so in the rare 8-cell stage embryos. We defined differentially expressed genes (DEGs) using thresholds of |log₂FC| > 1 and adjusted *P* < 0.05 across four pair-wise comparisons (Figure 1C; Table S2).

Volcano plots (Figure 1D) visualize the four key comparisons, and we provide the matching gene set enrichment analysis (GSEA) to each pair-wise contrast (Figure S2; Table S3): “To be normal” at the 2c (2cWT vs. 2cKO) and 8c (8cWT vs. 8cKO) stages; and “To develop” from 2c to 8c in WT (8cWT vs. 2cWT) and KO (8cKO vs. 2cKO) embryos. The volcano plots and an intersection analysis (Figures 1E–F) using a four-way Venn diagram approach revealed that most DEGs fell into the “To develop” category, consistent with the PCA results.

### Transcriptional programs of development do not collapse in the absence of maternal SETDB1

We first examined DEGs associated with the “To develop” category. We identified 1,722 genes downregulated during development in both WT (8cWT vs. 2cWT) and KO (8cKO vs. 2cKO) embryos (Figure 1E), suggesting their normal downregulation is not dependent on SETDB1. This trend was confirmed by heatmap clustering (Figure S3A).

Gene ontology (GO) terms associated with these downregulated DEGs included DNA binding, transcriptional activation, ubiquitination, and phospholipid binding (Figure S3B; Table S4). Similarly, 828 genes were consistently upregulated during development in both WT and KO embryos (Figure 1F) and were enriched in RNA production and translation (Figure S3D). Together, these results indicate that maternal SETDB1 is not required for the global execution of 2c-to-8c transcriptional transitions, and that the developmental arrest observed is not due to a complete transcriptional collapse.

### The two-cell program is misregulated in the absence of maternal SETDB1

To understand which developmental change occurs specifically in *Setdb1*^mat-/+^ embryos, we focused on genes differentially expressed only in KO embryos during the 2c to 8c transition. We identified 2,379 downregulated and 1,166 upregulated DEGs in the 8cKO vs. 2cKO contrast, but not in the WT developmental comparison (Figures 1E, 1F).

Heatmaps revealed that many genes exhibited aberrant expression in 2cKO embryos, with levels reverting to near-normal by the 8c stage (Figures S4A, S4C). Overrepresentation analysis indicated that 2cKO embryos underexpressed genes related to ribosomal and mitochondrial function (Figure S4B), while overexpressing genes associated with later developmental processes such as pattern specification and organ morphogenesis (Figure S4D). These observations suggest that mis-timed activation or repression occurs in the absence of maternal SETDB1.

In a related heat map comparison (Figure S4E), we identified genes uniquely overexpressed in 2cKO embryos that are enriched in ribosomal and mitochondrial pathways (Figure S4F). Thus, maternal SETDB1 is required to properly regulate 2-cell-stage gene expression dynamics.

### Maternal SETDB1 is required for timely suppression of cleavage-stage transcripts

To explore SETDB1’s role in establishing a normal 2c transcriptome, we analyzed 2cWT vs. 2cKO DEGs and found 171 downregulated and 238 upregulated transcripts (Figures 1E, 1F; Table S2). Among the 69 downregulated genes exclusive to 2cWT vs. 2cKO, many are prematurely activated in 2cKO embryos, including *Klf10*, *Lig1*, and *H1f3* (Figure 1G). Another 59 DEGs downregulated in 2cWT vs. 2cKO overlapped with 8cKO vs. 2cKO DEGs and were misexpressed only at the 2c stage (Figure 1H), suggesting these transcripts are normally repressed by SETDB1 at 2c but not at 8c. Notable example is *Zscan4c*, a key regulator of the 2-cell program in vitro [33].

To understand SETDB1’s role at the 8c stage, we analyzed 636 downregulated and 302 upregulated DEGs in 8cWT vs. 8cKO embryos (Table S2). Of these, 270 overlapped with 8cWT vs. 2cWT DEGs (Figure 1E), indicating that maternal SETDB1 normally suppresses these transcripts during the 2c to 8c transition (Figure 1I). This group included *Usp17ld*, *Tdpoz1* and *Tdpoz3*, transcripts expressed transiently at the 1-cell, 2-cell or 4-cell stages [34].

An additional 121 DEGs overlapped with both developmental and genotype comparisons, suggesting that their silencing occurs more slowly in *Setdb1*^mat-/+^ embryos than in controls (Figure 1J). These included the 2-cell transient genes [34] *Obox3* and *Fzd7*, further implicating maternal SETDB1 in regulating 2c-specific transcriptional shutdown. Taken together, this four-way comparison approach highlights maternal SETDB1 as a key factor for the timely suppression of specific transcripts at both the 2-cell and 8-cell stages.

### Maternal SETDB1 controls transient gene expression during ZGA

We next asked whether maternal SETDB1 regulates specific ZGA gene classes as defined in the “Database of Transcriptome in Mouse Early Embryos” (DBTMEE) [34]. Overrepresentation analysis (Figure 2A; Table S5) using DBTMEE gene clusters from Sakashita et al. [22] revealed distinct patterns across four two-way DEG comparisons.

**Figure 2.**
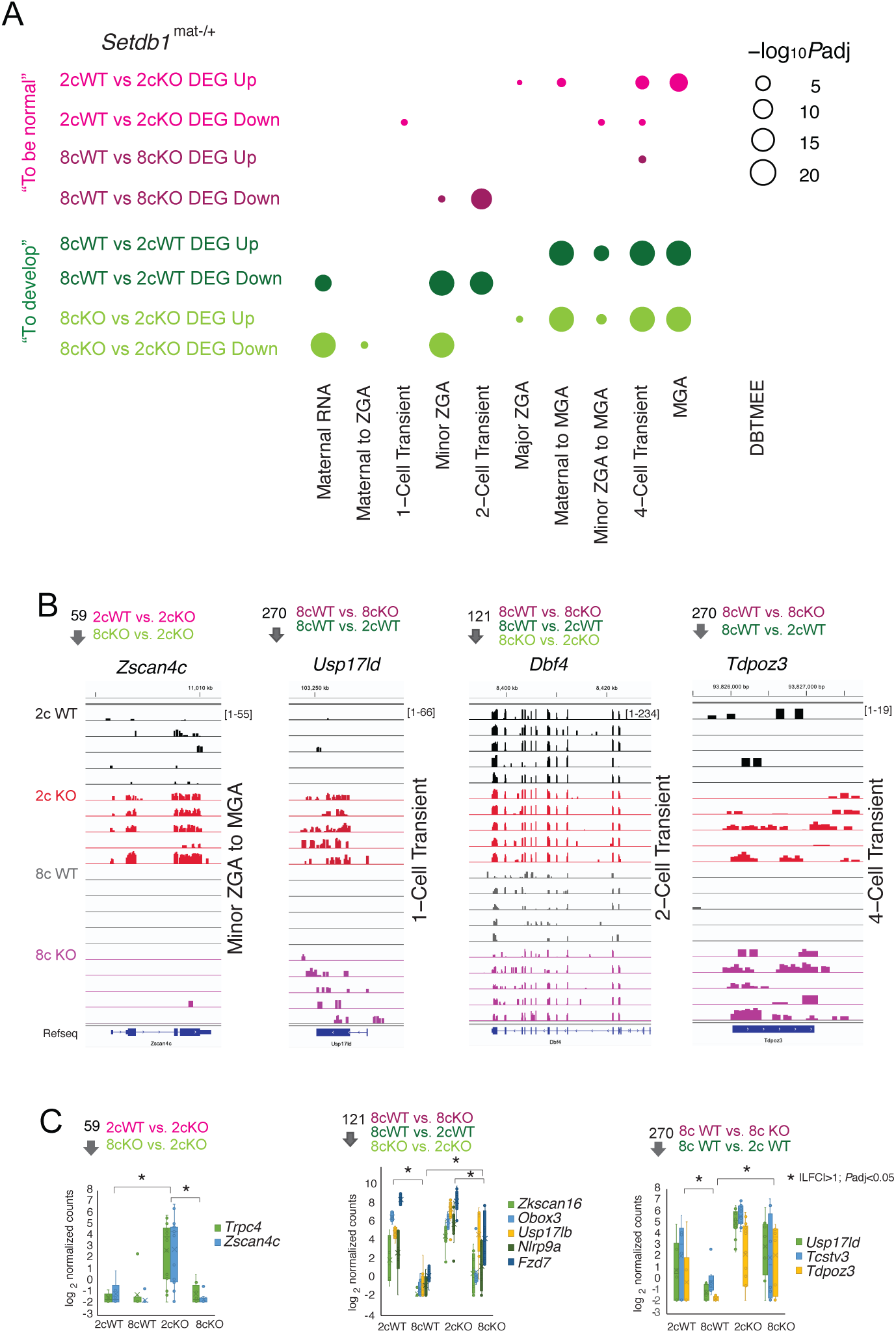
Maternal SETDB1 extinguishes 2-cell transient gene expression. (A) Bubble plot showing overrepresentation analysis of DBTMEE [34]-defined transcript sets among differentially expressed genes (DEGs) identified in the four pairwise comparisons. (B) IGV browser snapshots of representative transcripts from the DBTMEE-defined minor ZGA to MGA, 2-cell transient, and 4-cell transient gene sets across five biological replicates. Venn diagram compartments are indicated above each track. Unit in brackets represent normalized counts per million (CPM). Bigwig tracks are shown in the transcriptional direction matching the depicted gene. (C) Boxplots of selected DEGs *(|log₂FC| > 1, adjusted *P* < 0.05) from each Venn compartment, based on data from 2cWT (n=6), 2cKO (n=15), 8cWT (n=8), and 8cKO (n=8) embryos.

Maternal RNA and minor ZGA genes were appropriately downregulated in both WT and KO embryos from the 2c to 8c stage, as expected during normal development. However, 2-cell-specific transient transcripts were significantly downregulated in 8cWT vs. 2cWT embryos but not in 8cKO vs. 2cKO embryos. Moreover, 8cWT vs. 8cKO downregulated DEGs were enriched in this gene set, indicating a requirement for maternal SETDB1 to suppress 2c-transient transcripts by the 8c stage.

Similarly, 1-cell transient genes at the 2c stage and minor ZGA genes at the 8c stage were also insufficiently silenced in KO embryos. By contrast, mid-preimplantation gene activation (MGA) genes and their upstream regulators were upregulated normally in both genotypes. Notably, major ZGA and mid-preimplantation gene activity (MGA) transition-related genes were also underexpressed in 2cKO embryos, suggesting that SETDB1 may also support their proper induction.

Representative IGV browser tracks illustrate misregulation patterns in 5 biological replicates per condition (Figure 2B), including *Zscan4c* (minor ZGA), *Usp17ld* (*Dub1b*) (1-cell transient) *Dbf4* (2c transient), and *Tdpoz3* (4c transient). Differential expression status is mapped across Venn diagram segments (Figure 1E), and selected transcripts are shown in boxplots (Figure 2C).

These results demonstrate that maternal SETDB1 is required to suppress 1-cell transient genes by the 2c stage, and minor ZGA and 2c-specific genes by the 8c stage, ensuring transcriptional fidelity across cleavage stages.

### Maternal SETDB1 suppresses 2-cell-specific MERVL-chimeric transcripts

MERVL retroelements act as alternative promoters for chimeric transcripts during the 2-cell stage and serve as molecular hallmarks of totipotency [9]. Although SETDB1 represses ERVK and ERVL-MaLR retrotransposons in oocytes, it does not affect MERVLs at that stage [6].

We found significantly higher transcript levels of *MERVLint:ERVL:LTR* and *MT2_Mm:ERVL:LTR* in 2cKO than in 2cWT embryos (Figure 3A; Table S6). These transcripts were efficiently extinguished in WT embryos by the 8c stage but remained partially expressed in KO embryos. The difference was even more pronounced at 8c: *MT2* expression was downregulated in 8cWT vs. 8cKO with LFC = –2.65 and adj. *P* = 1.31E–16, compared to LFC = –1.86 and adj. *P* = 1.41E–08 in 2cWT vs. 2cKO.

**Figure 3.**
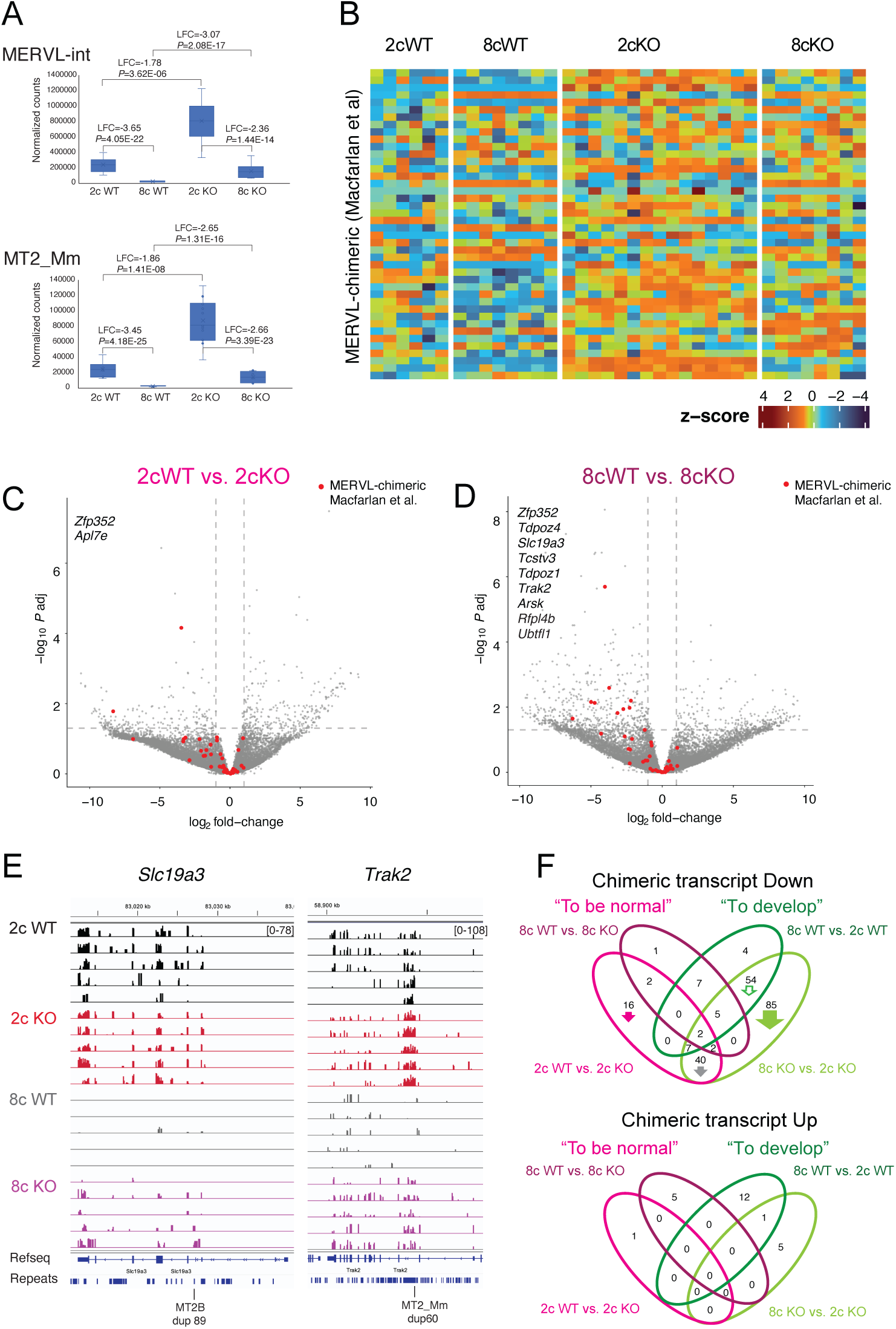
Maternal SETDB1 regulates MERVL-driven chimeric transcripts. (A) Boxplots showing normalized counts of multimapped MERVL-int and MT2_Mm elements. (B) Heatmap from *Setdb1* KO embryos at MERVL chimeric transcripts classified by Macfarlan [9]. (C–D) Volcano plots marking Macfarlan-defined chimeric transcripts in 2cWT vs. 2cKO and 8cWT vs. 8cKO pairwise comparisons. (E) IGV browser images of MT2B1 LTR-driven MERVL-chimeric transcripts. (F) Venn diagram showing differentially expressed (*P* < 0.05) known and novel TE-driven chimeric transcripts.

Similarly, many MERVL chimeric transcripts defined by Macfarlan et al. [9] were upregulated in both 2cKO and 8cKO embryos (Figure 3B), with most clustering on the left side of volcano plots (downregulated in WT vs. KO) (Figure 3C-D). Two shared DEGs reached significance at 2c, and nine at 8c (*P* adj < 0.05, |log₂FC| > 1). IGV views show misregulation of representative transcripts *Slc19a3* and *Trak2* (Figure 3E).

Using a combined list of previously identified [35] and novel chimeric transcripts we performed a four-way comparison to assess SETDB1 dependency (Figure 3F; Table S6). SETDB1 had a strong suppressive effect on TE-driven, including MERVL-driven transcripts at both 2c and 8c stages, with no clear activating role. Thus, maternal SETDB1 is essential to silence 2c-specific MERVL chimeric transcripts during preimplantation development.

### Maternal SETDB1 silences MERVL MT2 LTR-activated transcripts

MT2s represent major MERVL subfamilies and their LTRs can act as alternative promoters or distal enhancers, depending on their genomic position and orientation. A recent study by Yang et al. [23] used a CRISPR-based epigenetic MT2 LTR inactivation system (MT2i) to identify MT2-controlled genes in mouse embryos. Using ChIP-seq data from Wang et al. [26] we found that H3K9me3 deposition at MT2-dependent transcripts is first reduced between E2c and L2c but later increases from 4c to the 8c stage (Figure 4A). MT2s that drive those transcripts accumulate H3K9me3 peaks progressively from the E2c to 8c stages (Figure 4B), and these peaks are stronger and more spatially defined. Peaks localized at the MT2 edges is consistent with low internal mappability in repeat elements. IGV browser views show that the expression of MT2-controlled DEGs such as *Arsk* and *Usp17ld* was derepressed in 2cKO and 8cKO embryos while their flanking MERVLs and MT2s display strong H3K9me3 peaks in WT embryos (Figures 4C–E).

**Figure 4.**
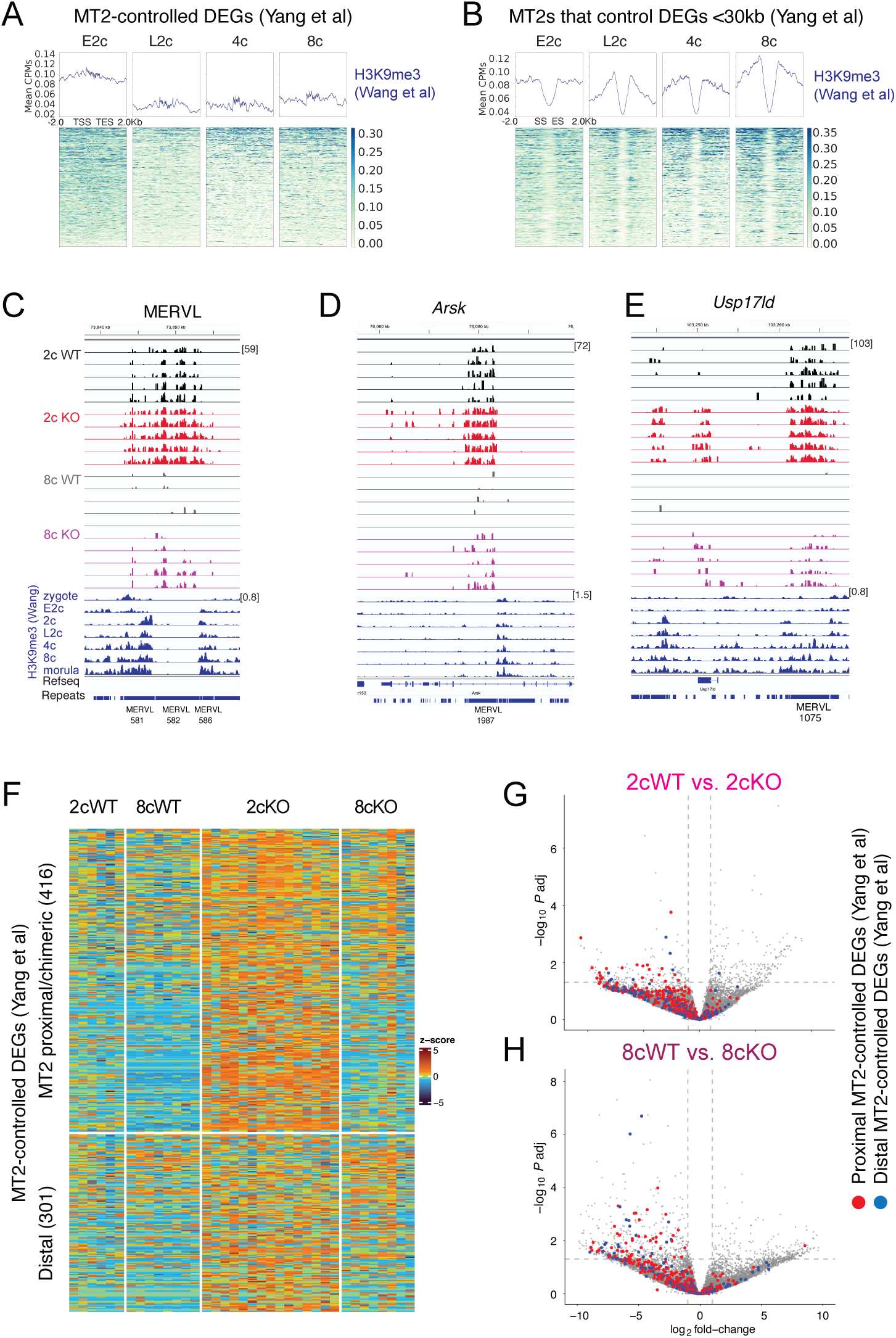
Maternal SETDB1 suppresses MT2 LTR-regulated genes. (A–B) Heatmaps of H3K9me3 deposition [26] at MT2-controlled [23] DEGs (A) and their MT2 elements (B) across early embryonic stages. (C–E) IGV browser views of a representative MERVL and MT2-regulated loci. (F) Heatmap of maternal *Setdb1* KO embryos at DEGs classified by MT2i data. (G–H) Volcano plots highlighting MT2-regulated genes in 2cWT vs. 2cKO and 8cWT vs. 8cKO pairwise comparisons.

Volcano plots reveal that the expression of MT2-controlled DEGs in general was derepressed in 2cKO and 8cKO embryos (Figures 4F–H). These genes clustered on the left of volcano plots, indicating strong SETDB1-mediated repression. Yang et al. [23] identified 341 genes requiring MT2 for activation by early 2c (E2c) and 1,111 by late 2c (L2c). We observed that many of these genes were derepressed in SETDB1 KO embryos (Figures 5A–E). These results provide genetic evidence that maternal SETDB1 is essential for silencing MT2-activated genes.

**Figure 5.**
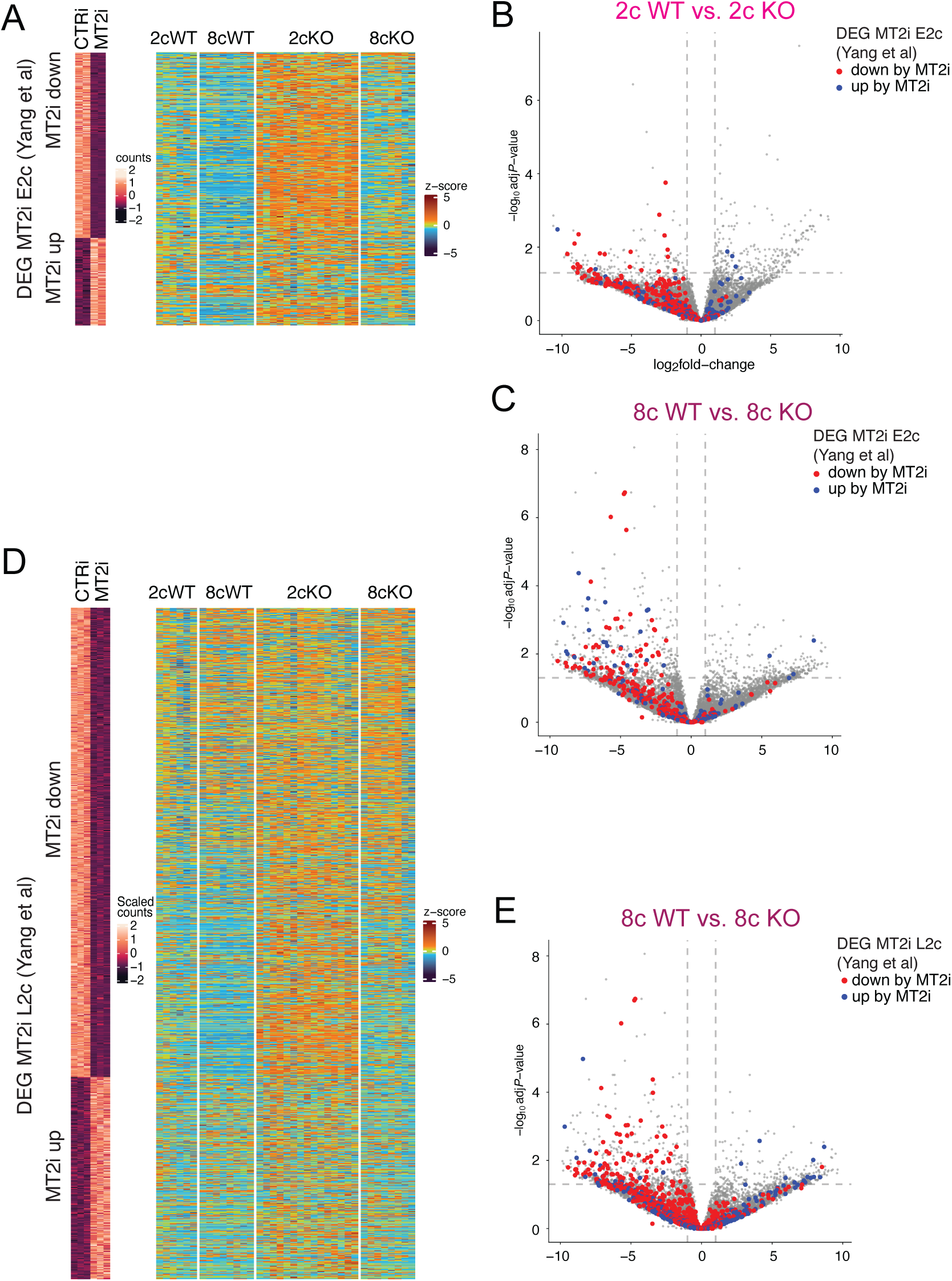
SETDB1 regulates MT2-activated genes across time. (A) Heatmaps comparing MT2i-responsive [23] early two cell (E2c) DEGs with the *Setdb1* KO transcriptomes. (B, C) Volcano plots highlighting MT2-regulated E2c DEGs in 2cWT vs. 2cKO and 8cWT vs. 8cKO pairwise comparisons. (D) Heatmaps of MT2i late two cell (L2c) DEGs. (E) Volcano plot highlighting MT2-regulated L2c DEGs.

### Maternal SETDB1 has a broad effect on transposable elements in 2c and 8c embryos

We next examined whether maternal SETDB1 broadly regulates TE expression during preimplantation. PCA of TE expression in our RNA-seq data revealed clustering first by stage and then by genotype, with increased variability among 8cKO samples (Figure S5A). We analyzed both multimapped and uniquely mapped TE reads. Multimapped TE analysis (Figure S5B; Table S7) showed that maternal SETDB1 is required to suppress TEs in WT vs. KO embryos and plays a role during 2c to 8c development.

The tallies of “To be normal” DE TEs by TE families show that several TE families— including ERVK, ERVL, ERVL-MaLR, ERV1, and L1 elements—require suppression by SETDB1 in WT embryos (Figure S5C). The ERVK, ERVL-MaLR, and ERV1 families were most affected in both 2cWT vs. 2cKO and 8cWT vs. 8cKO contrasts, while LINEs required SETDB1 for repression mainly at 2c. The tallies of “To develop” suggest that SETDB also affects developmental TE changes during 2c-to-8c transitions. ERVK and LINE-1 elements appear to be developmentally activated in WT but less so in KO embryos (Figure S5D), which we investigate further below.

Heatmaps of “To be normal” DE TEs (Figure S5E) revealed strong derepression of nearly all repeat families in KO embryos at both the 2c and 8c stages. Heatmaps of “To develop” DE TEs (Figure S5F) clarify how developmental changes are affected by SETDB1. In WT embryos, L1 LINEs and many ERVKs were upregulated between 2c and 8c, while MERVLs and ERVL-MaLRs were appropriately silenced. This developmental regulation was disrupted in KO embryos: 8cKO vs. 2cKO showed partial MERVL/ERV1 repression and inappropriate activation of some ERVKs and LINEs—due to the highly derepressed state in 2cKO embryos. The analysis of uniquely mapped TE transcripts in presented in Table S8.

These results collectively show that maternal SETDB1 is a major regulator of TE expression, with broad effects extending beyond MERVLs to multiple TE families during early development.

### Maternal SETDB1 is required to extinguish DUXBL-responsive transcripts

DUXBL is a known repressor of ZGA and MERVL genes, acting in the footsteps of DUX during the exit from totipotency [25]. We tested whether maternal SETDB1 is required for DUXBL-mediated suppression. We compared our RNA-seq data to the top 50 DEGs from *Duxbl*^-/-^ embryos [25] and found that 7 and 12 DUXBL-suppressed transcripts were derepressed in 2cKO, and 8cKO embryos, respectively (Figure 6A), suggesting that maternal SETDB1 acts specifically in their silencing. Volcano plots (Figure 6B) showed DUXBL-upregulated genes clustering on the left in both 2cWT vs. 2cKO and 8cWT vs. 8cKO comparisons, indicating again SETDB1-dependent suppression.

**Figure 6.**
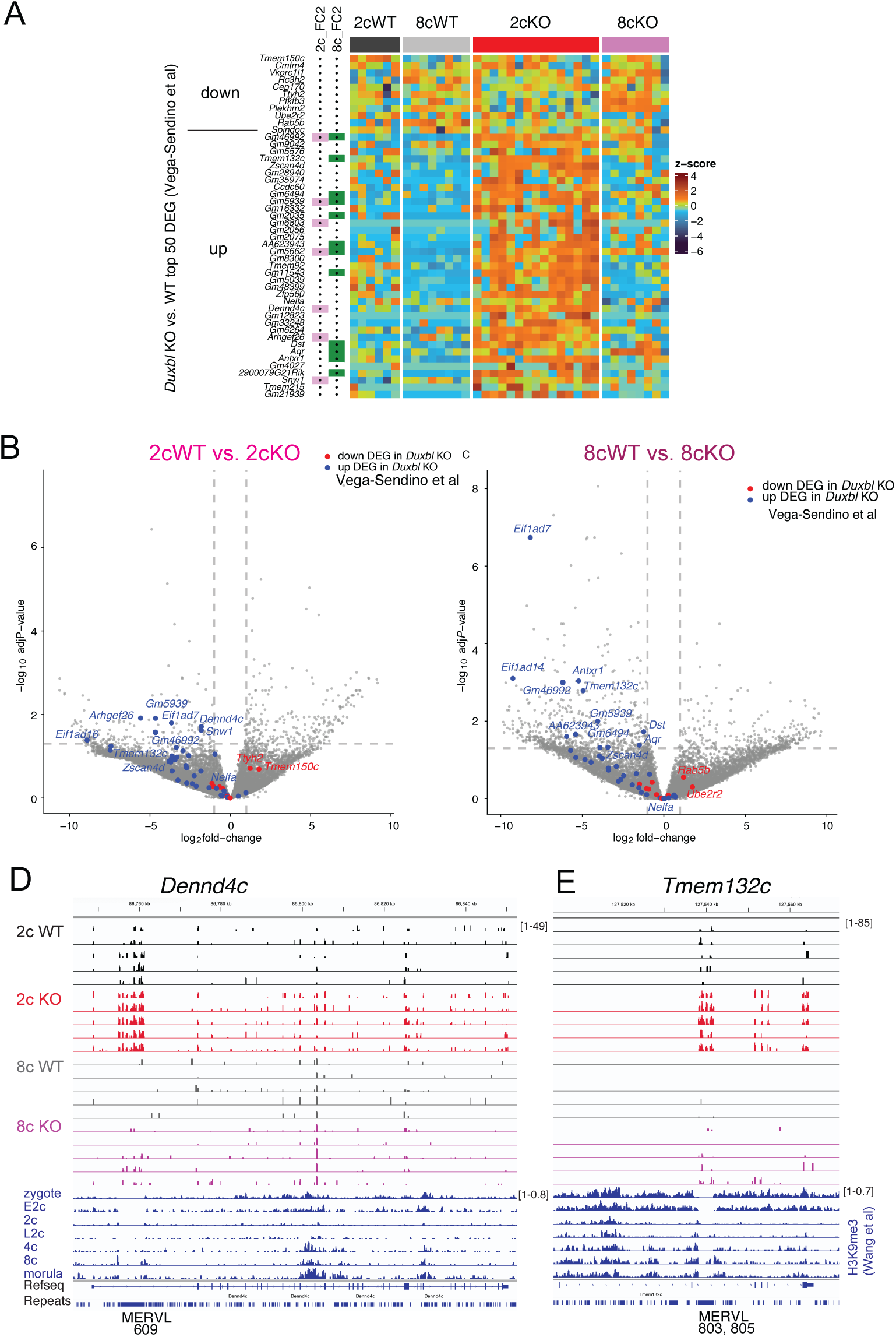
SETDB1 represses DUXBL-responsive transcripts. (A) Heatmap of top 50 DEGs identified in *Duxbl* KO embryos [25] analyzed in the *Setdb1* KO RNA-seq dataset. (B) Volcano plots marking up-/downregulated DUXBL targets in 2cWT vs. 2cKO and 8cWT vs. 8cKO pairwise comparisons. (D–E) IGV browser examples of DUXBL-responsive DEGs with aligned H3K9me3 ChIP-seq data.

IGV browser views confirmed that *Dennd4c* (2c), *Tmem132c*, *Antxr1*, and *Aqr* (8c) were derepressed in KO embryos and associated with H3K9me3 peaks flanking MERVL elements in WT embryos (Figures 6D–E; S6A–B). Although H3K9me3 signal within MERVLs is low due to mappability limits, adjacent flanks exhibited strong enrichment beginning at the 2c stage and increasing through 4c to morula Wang et al. [26] Two additional genes, *Nelfa* and *Zscan4d*, appeared derepressed in KO embryos (Figure S6C–D), despite not crossing significance thresholds. MERVLs serve as alternative promoters or distal enhancers for these transcripts, with H3K9me3 deposited near these elements in WT but not KO embryos.

Together, these data provide genetic evidence that DUXBL requires maternal SETDB1 to silence a subset of its targets via H3K9me3 deposition at MERVL LTRs.

### Overrepresentation analysis confirms shared targets with MT2 and DUXBL pathways

We performed ORA to test for overlap between SETDB1-sensitive genes and those reported by Yang et al. (MT2i) [23] and Vega-Sendino et al. (DUXBL KO) [25] (Figure 7A; Table S9). We found highly significant overlap between: Proximal MT2-controlled DEGs and DEGs downregulated in 2cWT vs. 2cKO (adj. *P* = 3.19E–04); Proximal MT2-controlled DEGs and DEGs in 8cWT vs. 8cKO (adj. *P* = 7.17E–04); E2c MT2i-downregulated genes and SETDB1-repressed DEGs at 2c (adj. *P* = 8.18E–05); and E2c MT2i-downregulated genes and SETDB1-repressed DEGs at 8c (adj. *P* = 5.77E–11). This suggests that genes activated by MT2s during ZGA also require maternal SETDB1 for their timely silencing later.

**Figure 7.**
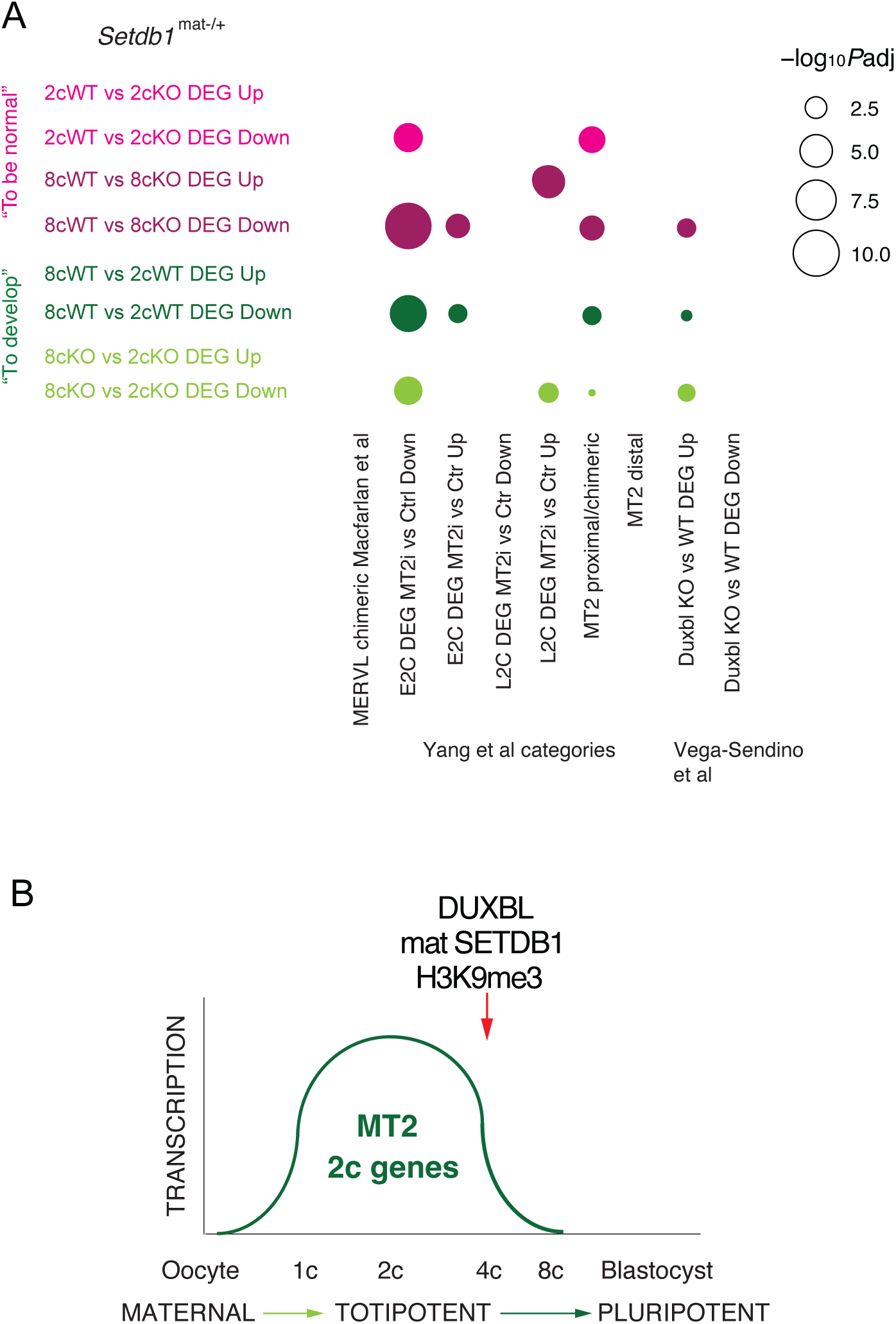
SETDB1 collaborates with DUXBL to repress totipotency programs. (A) Bubble plot of overrepresentation analysis showing enrichment of previously identified MERVL-chimeric, MT2i-responsive and *Duxbl* KO-responsive gene sets [9, 23, 25] in *Setdb1* KO pairwise DEGs. (B) Summary model: Maternal SETDB1 deposits H3K9me3 at MT2 elements to silence MERVL-driven 2-cell transcripts in coordination with DUXBL, enabling exit from totipotency.

We also observed strong enrichment between DUXBL-suppressed genes and SETDB1-repressed DEGs in 8cWT vs. 8cKO embryos (*P* adj = 0.0071), supporting a cooperative role in extinguishing the 2c program.

In summary, these findings establish SETDB1, a H3K9 methyltransferase, as a broad-acting chromatin regulator in the preimplantation-stage embryo required for: silencing MERVL retroviral elements; extinguishing minor ZGA and transient totipotency-associated genes; and terminating transcription of MT2-activated and DUXBL-regulated transcripts at key regulatory elements in cleavage-stage embryos.

## Discussion

SETDB1 is essential for mammalian development due to its role in establishing transcriptionally repressive H3K9me3 marks. Our findings indicate that the maternal-effect lethality observed in *Setdb1*^mat-/+^ preimplantation embryos is not due to a global transcriptional collapse, but rather the failure to silence specific transcriptional programs that must be extinguished for normal progression through, and beyond, the cleavage stages. Exit from the totipotency state is essential for embryonic development [18, 24]. While SETDB1 affects a broad range of genes and retroelements during preimplantation development, its specific role in silencing the totipotency program alone underscores its critical role as a chromatin regulator in early embryogenesis.

Maternal SETDB1 functions in a highly specific and dynamic manner. At the 2-cell stage, it suppresses 1-cell transient and minor ZGA transcripts; by the 8-cell stage, it silences 2-cell transient and MERVL-associated chimeric transcripts. In normal development, these silencing events coincide with dynamic H3K9me3 deposition at MERVLs and MT2 LTRs. In ESCs, SETDB1 prevents dedifferentiation into 2-cell-like cells by depositing H3K9me3 at or near totipotency genes [31], suggesting that a similar mechanism may operate during embryo development in vivo. However, the rarity of *Setdb1*^mat-/+^ 8-cell embryos limits our ability to directly confirm H3K9me3 depletion at these loci—a limitation of this study.

SETDB1’s mechanism likely involves recruitment to specific retroelement loci, such as MT2 LTRs by transcription factors, such as DUXBL. Our genetic data show that SETDB1 is required for silencing both MT2 LTRs and DUXBL-responsive transcripts. The overlap of SETDB1-regulated genes with those affected by MT2 inactivation [23] and DUXBL loss [25], along with the presence of H3K9me3 at these loci during normal preimplantation development, suggests a coordinated silencing mechanism in the embryo. Phenotypic parallels between *Setdb1*^mat-/+^ and *Duxbl*^-/-^ embryos support this model. While *Duxbl*^-/-^ embryos arrest at the 4-cell stage [25], a few *Setdb1*^mat-/+^ embryos progress to the 8-cell stage, suggesting that SETDB1 may act redundantly with, or downstream of, DUXBL.

In conclusion, we identify maternal SETDB1 as a critical in vivo epigenetic regulator that extinguishes totipotency-associated transcriptional programs, particularly those driven by retroelements and responsive to DUXBL. Its role at MT2 elements supports a model in which SETDB1 provides an epigenetic memory, silencing retroelements shortly after their activation by DUX and OBOX. This mechanism rapidly restricts the 2-cell transcriptional program, enabling the transition to pluripotency. By repressing MT2 LTRs, SETDB1 ensures successful progression beyond the cleavage stages and supports the earliest cell fate transitions in the mammalian embryo (Figure 7B).

## Materials and Methods

### Mice

All animal experiments were conducted in accordance with the National Research Council’s Guide for the Care and Use of Laboratory Animals, under Institutional Animal Care and Use Committee-approved protocols at Van Andel Institute (VAI). The *Setdb1* conditional knockout mouse line (*Setdb1*^f/f^) was derived from *Setdb1*^tm1a(EUCOMM)Wtsi^ (European Mouse Mutant Archive) as previously described [32]. The *Pgk*-neo cassette was removed via FLPE-mediated recombination using B6.Cg-Tg^(Pgk1-flpo)10Sykr/J^ (RRID:IMSR_JAX:011065) [36]. *Zp3*-cre transgenic mice C57BL/6-Tg^(Zp3-cre)93Knw/J^ [37] were used to excise SET-domain–encoding exons in growing oocytes. *Setdb1*^f/f^; *Zp3*-cre females were experimental animals; *Setdb1*^f/f^ females without the transgene served as controls.

### Collection of preimplantation mouse embryos

Embryos were collected at late 2-cell and 8-cell stages. For 2-cell embryos, 6–8-week-old females were superovulated with 5 IU PMSG followed by 5 IU hCG after 46–48 h, then mated with WT males. For 8-cell embryos, CARD HyperOva was used in 26–30-day-old females to increase efficiency. Embryos were harvested in M2 medium at 44 h (1.5 dpc, 2-cell) and 68 h (2.5 dpc, 8-cell) post-hCG. Embryos were washed in PBS, placed into 1.5 mL tubes, lysed in RNA-Bee, flash frozen, and stored at –80 °C.

### RNA isolation

RNA was isolated from single embryos using RNA-Bee. Linear polyacrylamide (LPA) was used for carrier-mediated isopropanol precipitation. RNasin® (Promega) was added during resuspension. DNA contamination was removed using rDNase I (Ambion). RNA was recovered after DNase inactivation and ethanol precipitation and eluted in RNasin-containing Tris buffer.

### Library preparation and sequencing

Libraries were prepared with the SMART-Seq Stranded Kit (Takara). Samples were randomized and processed with fragmentation to 250 bp (85 °C, 10 min embryo RNA or 6 min control RNA). Unique dual indexes were added during PCR1 (10 cycles), and 8 samples were pooled for PCR2 (12 cycles). Libraries were QC’d using Agilent chips, QuantiFluor®, and KAPA qPCR, and sequenced on a NovaSeq S2 (paired-end 50 bp, ∼25M reads/library).

### RNA-seq analysis

Reads were trimmed using Trim Galore (v0.60) [38, 39] and aligned to mm10 with STAR (v2.7.8) [40] using --quantMode GeneCounts. DGE analysis was performed with edgeR (v4.4.1) [41–43], using Benjamini-Hochberg adjustment (adjusted *P*<0.05). Normalized log_2_ counts were used in figures. Bigwig tracks were created using bamCoverage (deepTools v3.5.2, CPM normalization) [44]. Gene set enrichment analysis (GSEA) and overrepresentation analysis (ORA) against gene ontology (GO) and custom gene sets were performed with ClusterProfiler (v4.14.4) (RRID:SCR_016884) [45].

### Transposable element (TE) analysis

Two sets of data were analyzed for TE: multimapped reads and uni-mapped reads. For multimapped reads, STAR (v2.7.8) [40] was run with --winAnchorMultimapNmax 200, -- outFilterMultimapNmax 100, and --outSAMmultNmax 1. TEtranscripts (v2.2.3) [46] was used for multi-mapped TE counts. Uni-mapped reads were filtered with -q 255 using Samtools (v1.17) (RRID:SCR_002105) [47]. FeatureCounts (Subread v2.0.0) [48] was used with mm10_rmsk_TE.gtf to generate count matrices Differential expression was assessed using DESeq2 (v1.46.0) [49].

### Chimeric transcript analysis

Chimeric transcripts were quantified with Retrotransposon (v37), which uses STAR [48] and StringTie2 [50]. Count tables were processed using DESeq2 (RRID:SCR_000154) [49] for DGE (adjusted *P*<0.05).

### Public datasets

ChIP-seq fastq files [26] (GSE98149) were trimmed using Trim Galore (v0.60) [38, 39], aligned to mm10 with BWA (v0.7.1) (RRID:SCR_010910) [51], and filtered with Samtools (v1.17) (RRID:SCR_002105) [47]. BAMs were converted to BigWig and visualized with deepTools [44]. Supplementary data was used from other studies [5, 9, 23, 25, 34], for comparative analyses.

### Data visualization

Heatmaps were generated with ComplexHeatmap (R) [52]., bubble plots were generated using ggplot2 [53], and Venn diagrams using ggvenn (v0.1.10) [54].

## Data Availability

All data are available in the main text or the supplementary materials except the deep sequencing data, which have been deposited to GEO under accession GSE269417.

## Acknowledgments

We gratefully acknowledge the support from the Van Andel Institute (VAI) Genomics Core (RRID: SCR_022913), the VAI Bioinformatics and Biostatistics Core (RRID: SCR_024762) and the VAI Vivarium (RRID:SCR_023211). This work was supported by the Van Andel Institute.

## Conflict of interest disclosure

Authors declare that they have no competing interests.

## Supplementary Data

Supplemental Figures S1 to S6

Table S1

Legends to Supplemental Table S2 to S8 (separate files)

**Figure S1.**
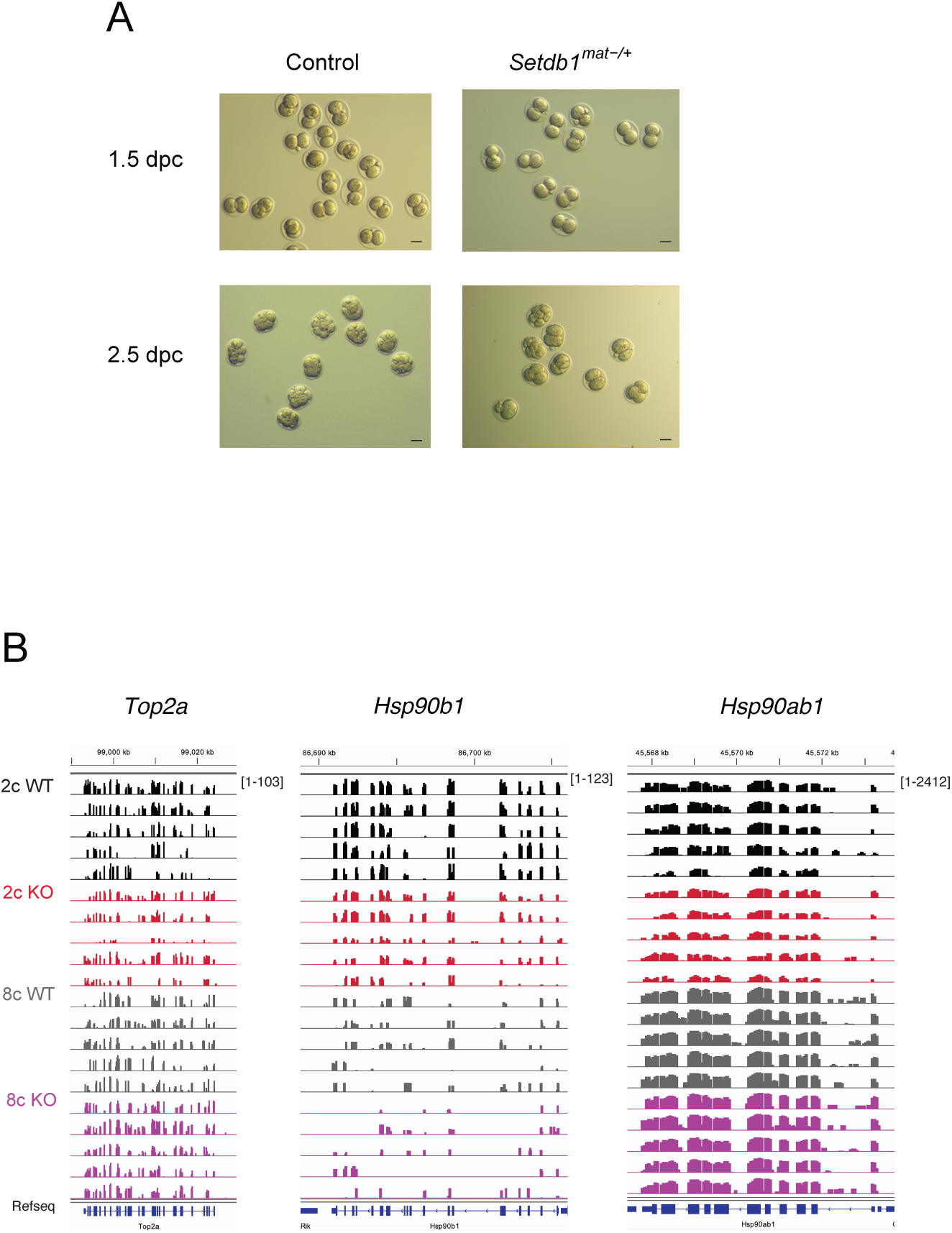
Normal features of *Setdb1* KO embryos. (A) Morphology. Brightfield images of 2c and 8c WT and KO embryos at 1.5 and 2.5 dpc. Scale bar 50 µM. (B) Transcription. Single embryo total RNA sequencing results of 2c and 8c WT and KO embryos at 1.5 and 2.5 dpc (n=5). IGV browser views of selected transcripts, with normalized CPM scales shown in brackets.

**Figure S2.**
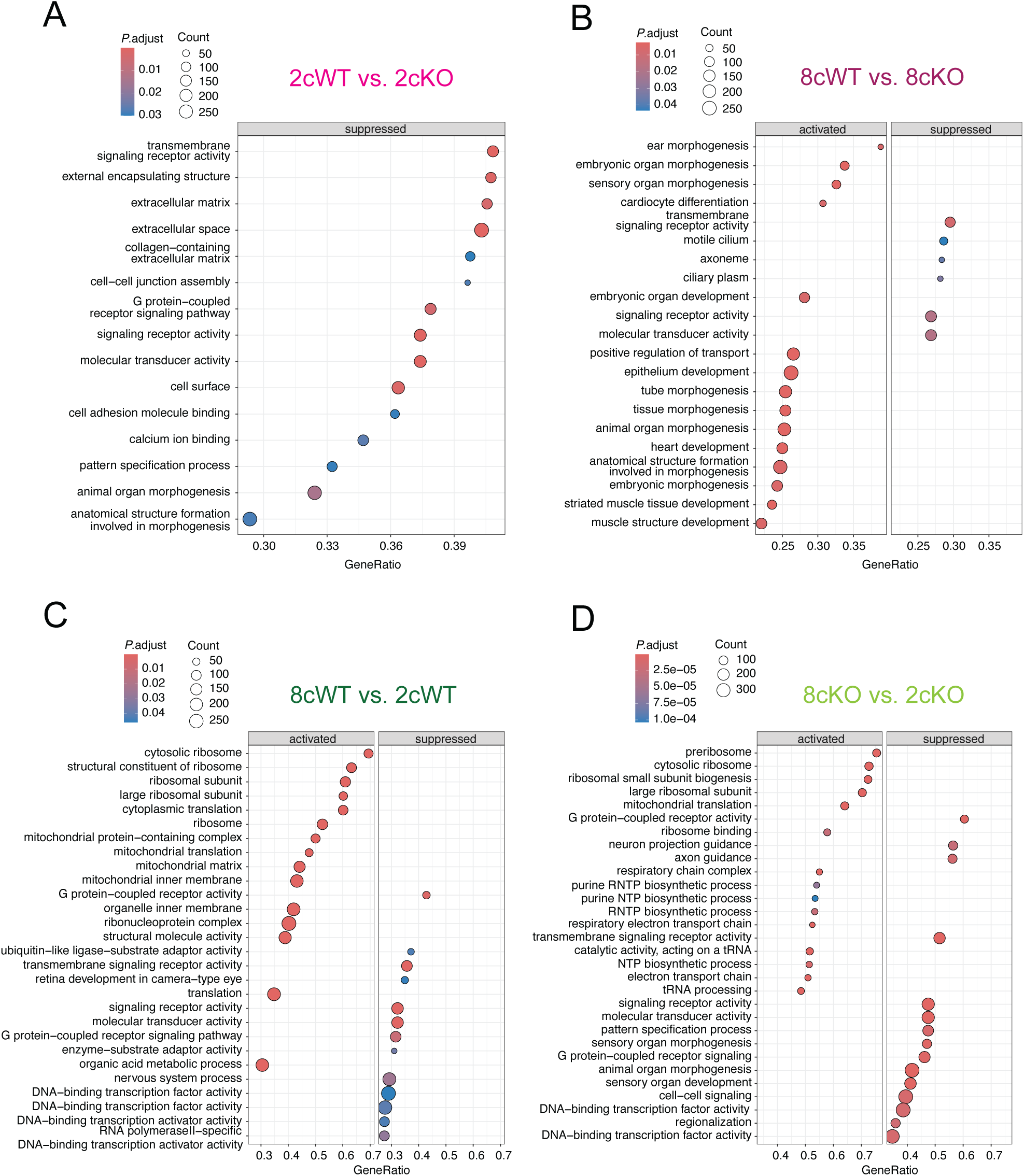
Gene set enrichment analyis (GSEA) of *Setdb1* KO DEGs. (A–D) Bubble plots of GSEA for 2cWT vs. 2cKO, 8cWT vs. 8cKO, 8cWT vs. 2cWT, and 8cKO vs. 2cKO pairwise comparisons.

**Figure S3.**
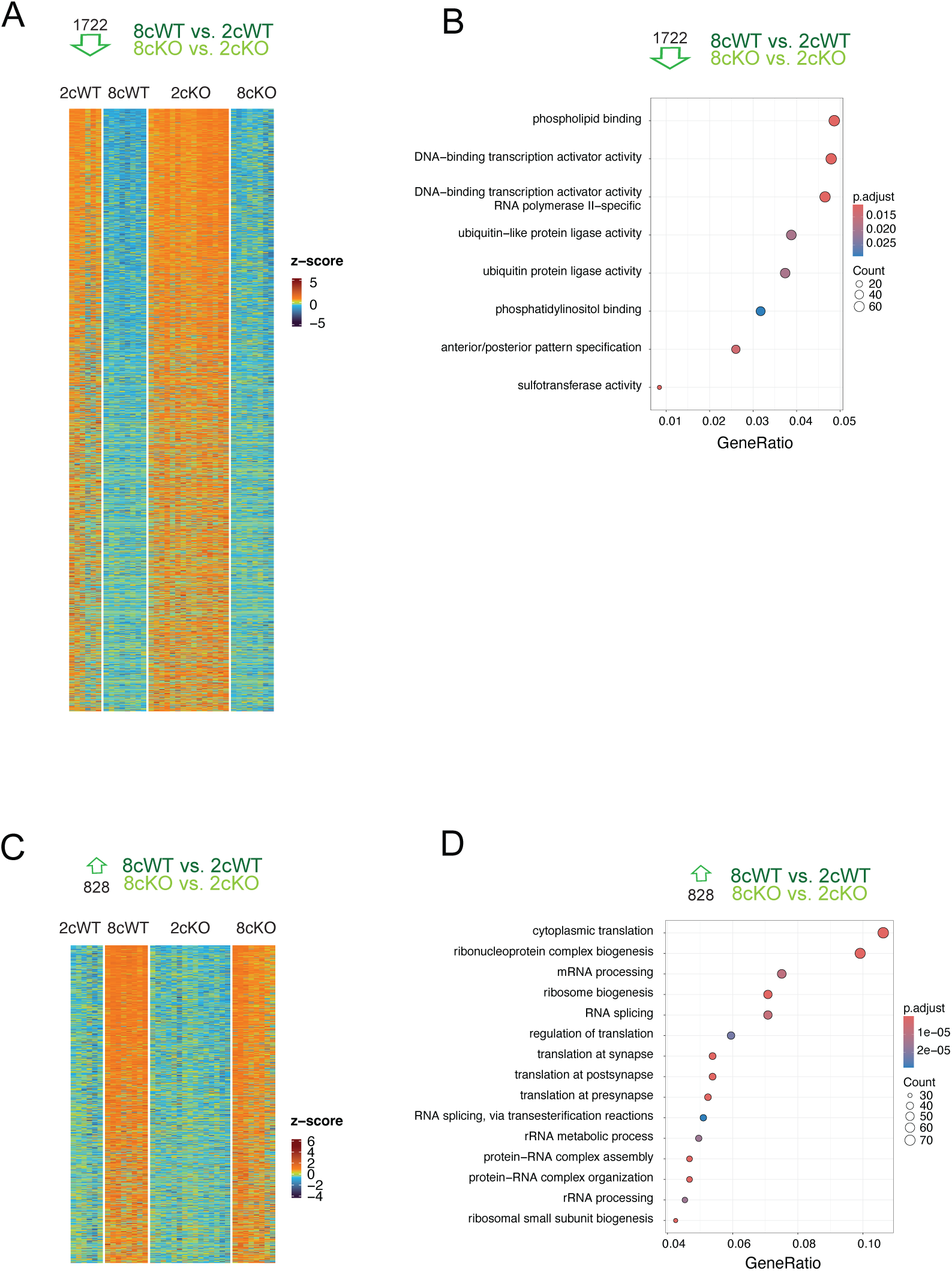
Developmental transcriptional changes independent of maternal SETDB1. (A, C) Heatmaps of DEGs commonly downregulated (n=1722) or upregulated (n=828) from 2c to 8c stages. (B, D) GO analysis of those DEGs.

**Figure S4.**
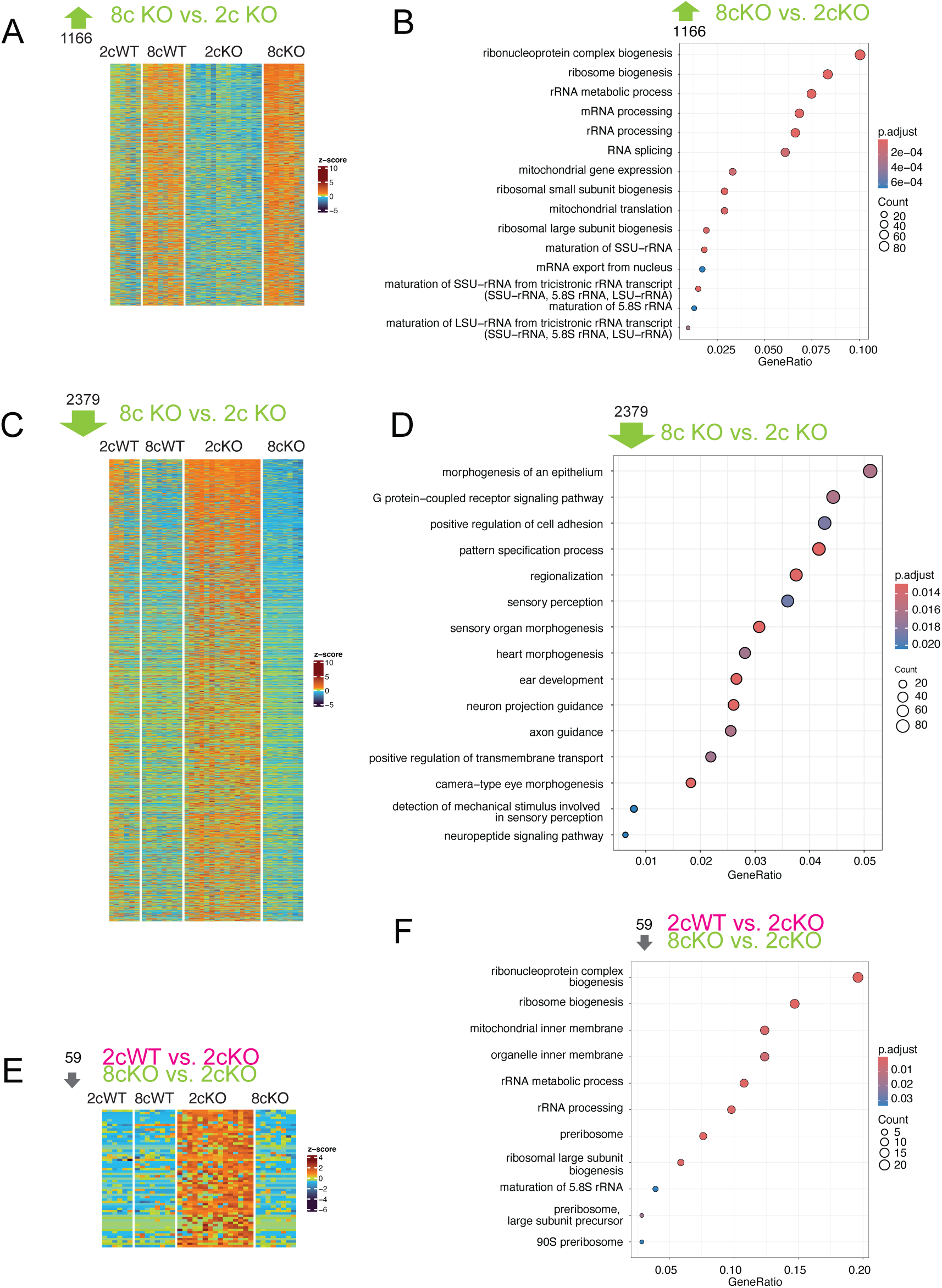
Developmental misregulation in *Setdb1* KO embryos. (A, C, E) Heatmaps showing DEGs uniquely changed in KO embryos. (B, D, F) GO enrichment analyses of these misregulated gene sets.

**Figure S5.**
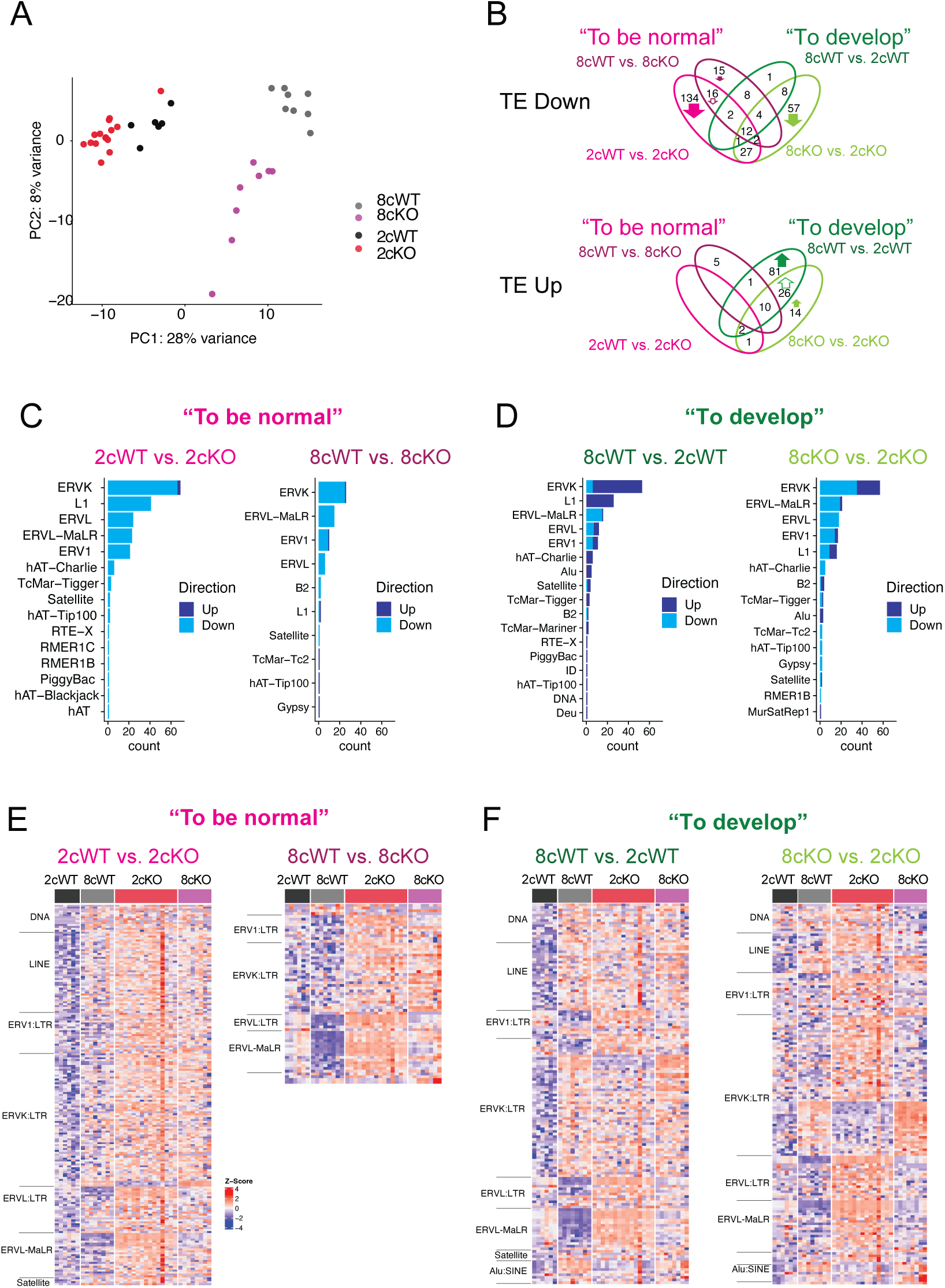
SETDB1 suppresses transposable elements at cleavage stages. (A) PCA of multimapped TE profiles. (B) Venn diagram of DE TE families (adjusted *P* < 0.05). (C–D) DE TE tallies across pairwise comparisons by TE class. (E-F) DE TE heatmaps across pairwise comparisons by TE class.

**Figure S6.**
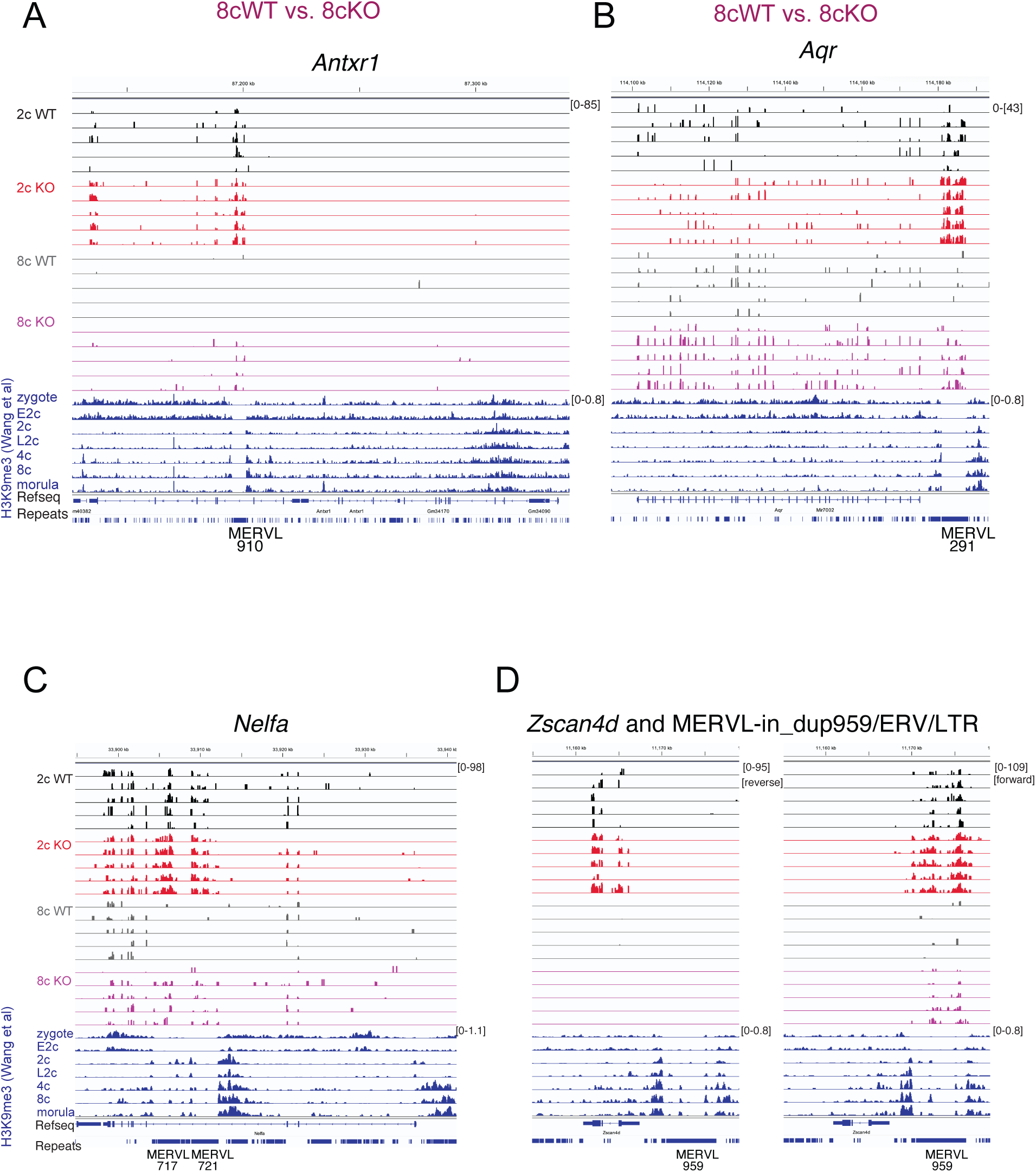
Examples of DUXBL targets derepressed in *Setdb1* KO embryos. (A–D) IGV images of *Duxbl* KO DEGs (e.g., *Antxr1*, *Aqr*, *Nelfa*, *Zscan4d*) with associated H3K9me3 profiles. MERVL insertions upstream or intronic are indicated.

**Table S1.** Sample Information. Details of all single embryos analyzed in the study, including genotype, developmental stage, and sequencing sample ID.

| **ID** | **Genotype** | **Stage** | **Collection**<br>**Day post**<br>**Coitum** | **Collection**<br>**Time** | **Name** |
| --- | --- | --- | --- | --- | --- |
| TZSC93 | Setdb1_MATKO | 2c | 1.5 | 10:00 AM | 2c KO |
| TZSC94 | Setdb1_MATKO | 2c | 1.5 | 10:00 AM | 2c KO |
| TZSC95 | Setdb1_MATKO | 2c | 1.5 | 10:00 AM | 2c KO |
| TZSC96 | Setdb1_MATKO | 2c | 1.5 | 10:00 AM | 2c KO |
| TZSC97 | Setdb1_MATKO | 2c | 1.5 | 10:00 AM | 2c KO |
| TZSC98 | Setdb1_MATKO | 2c | 1.5 | 10:00 AM | 2c KO |
| TZSC99 | Setdb1_MATKO | 2c | 1.5 | 10:00 AM | 2c KO |
| TZSC100 | Setdb1_MATKO | 2c | 1.5 | 10:00 AM | 2c KO |
| TZSC101 | Setdb1_MATKO | 2c | 1.5 | 10:00 AM | 2c KO |
| TZSC102 | Setdb1_MATKO | 2c | 1.5 | 10:00 AM | 2c KO |
| TZSC103 | Setdb1_MATKO | 2c | 1.5 | 10:00 AM | 2c KO |
| TZSC104 | Setdb1_MATKO | 2c | 1.5 | 10:00 AM | 2c KO |
| TZSC105 | Setdb1_MATKO | 2c | 1.5 | 10:00 AM | 2c KO |
| TZSC106 | Setdb1_MATKO | 2c | 1.5 | 10:00 AM | 2c KO |
| TZSC107 | Setdb1_MATKO | 2c | 1.5 | 10:00 AM | 2c KO |
| TZSC108 | Setdb1_MATKO | 2c | 1.5 | 10:00 AM | 2c KO |
| TZSC109 | Setdb1_MATKO | 8c | 2.5 | 10:00 AM | 8c KO |
| TZSC110 | Setdb1_MATKO | 8c | 2.5 | 10:00 AM | 8c KO |
| TZSC111 | Setdb1_MATKO | 8c | 2.5 | 10:00 AM | 8c KO |
| TZSC112 | Setdb1_MATKO | 8c | 2.5 | 10:00 AM | 8c KO |
| TZSC113 | Setdb1_MATKO | 8c | 2.5 | 10:00 AM | 8c KO |
| TZSC114 | Setdb1_MATKO | 8c | 2.5 | 10:00 AM | 8c KO |
| TZSC115 | Setdb1_MATKO | 8c | 2.5 | 10:00 AM | 8c KO |
| TZSC116 | Setdb1_MATKO | 8c | 2.5 | 10:00 AM | 8c KO |
| TZSC117 | Setdb1_WT | 2c | 1.5 | 10:00 AM | 2c WT |
| TZSC118 | Setdb1_WT | 2c | 1.5 | 10:00 AM | 2c WT |
| TZSC119 | Setdb1_WT | 2c | 1.5 | 10:00 AM | 2c WT |
| TZSC120 | Setdb1_WT | 2c | 1.5 | 10:00 AM | 2c WT |
| TZSC121 | Setdb1_WT | 2c | 1.5 | 10:00 AM | 2c WT |
| TZSC122 | Setdb1_WT | 2c | 1.5 | 10:00 AM | 2c WT |
| TZSC123 | Setdb1_WT | 2c | 1.5 | 10:00 AM | 2c WT |
| TZSC124 | Setdb1_WT | 2c | 1.5 | 10:00 AM | 2c WT |
| TZSC125 | Setdb1_WT | 8c | 2.5 | 10:00 AM | 8c WT |
| TZSC126 | Setdb1_WT | 8c | 2.5 | 10:00 AM | 8c WT |
| TZSC127 | Setdb1_WT | 8c | 2.5 | 10:00 AM | 8c WT |
| TZSC128 | Setdb1_WT | 8c | 2.5 | 10:00 AM | 8c WT |
| TZSC129 | Setdb1_WT | 8c | 2.5 | 10:00 AM | 8c WT |
| TZSC130 | Setdb1_WT | 8c | 2.5 | 10:00 AM | 8c WT |
| TZSC131 | Setdb1_WT | 8c | 2.5 | 10:00 AM | 8c WT |
| TZSC132 | Setdb1_WT | 8c | 2.5 | 10:00 AM | 8c WT |

## Supplemental Tables S2-S9. (separate files)

**Table S2.**
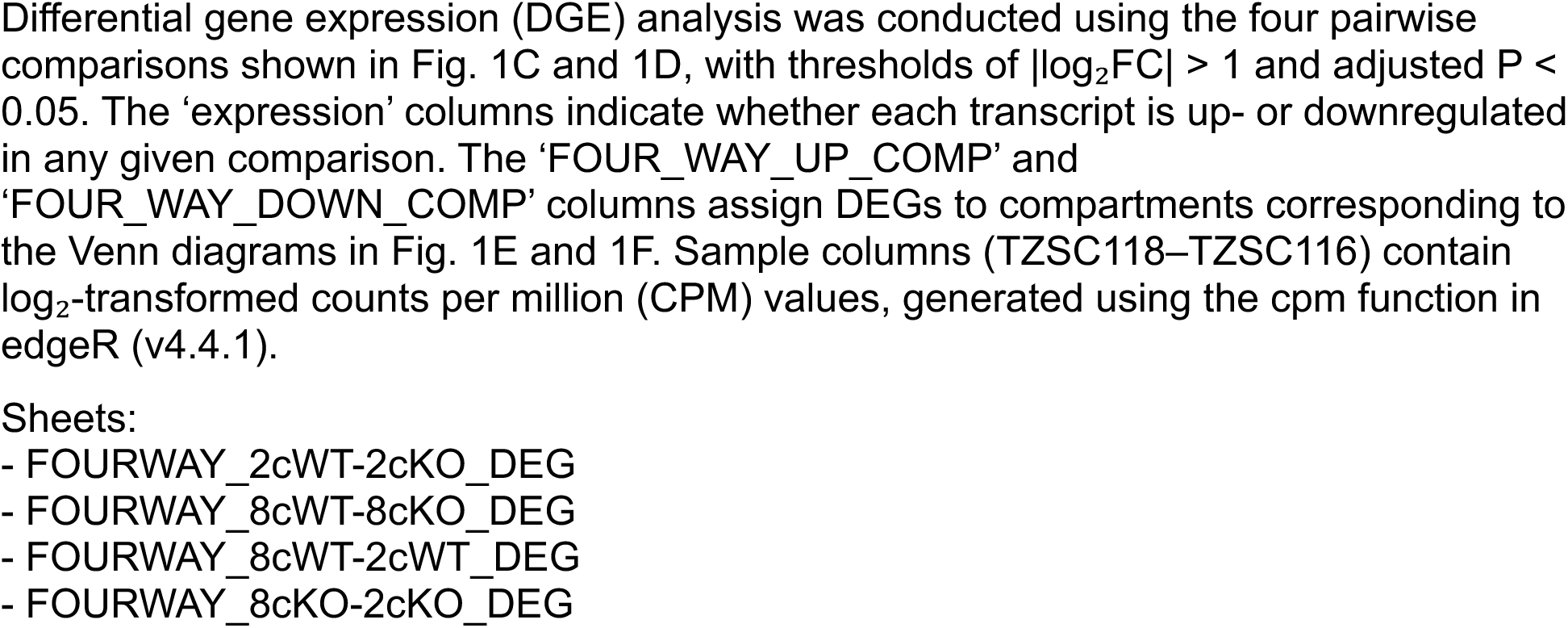
Differential Gene Expression (DGE) Analysis.

**Table S3.**
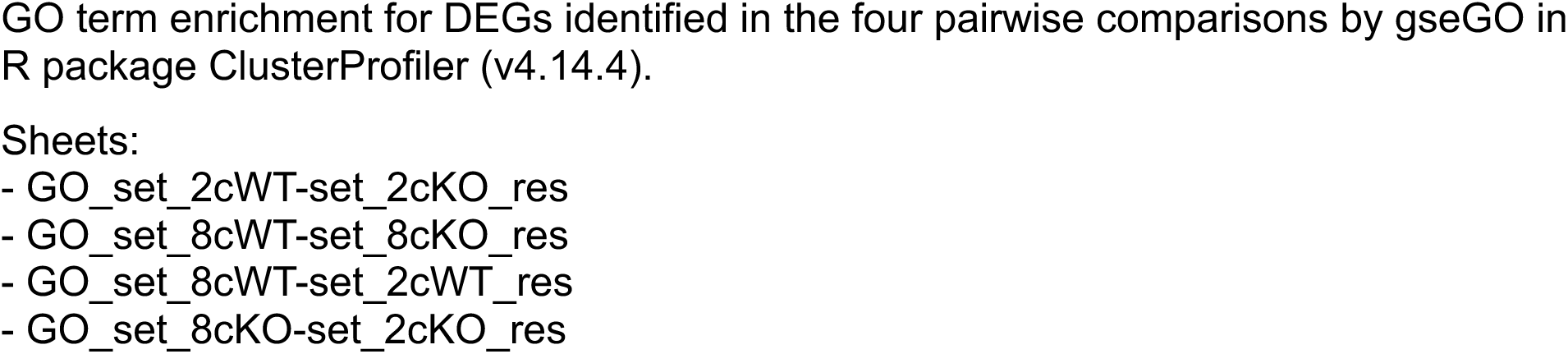
Gene Set Enrichment Analysis (GSEA) on Gene ontology (GO.

**Table S4.**
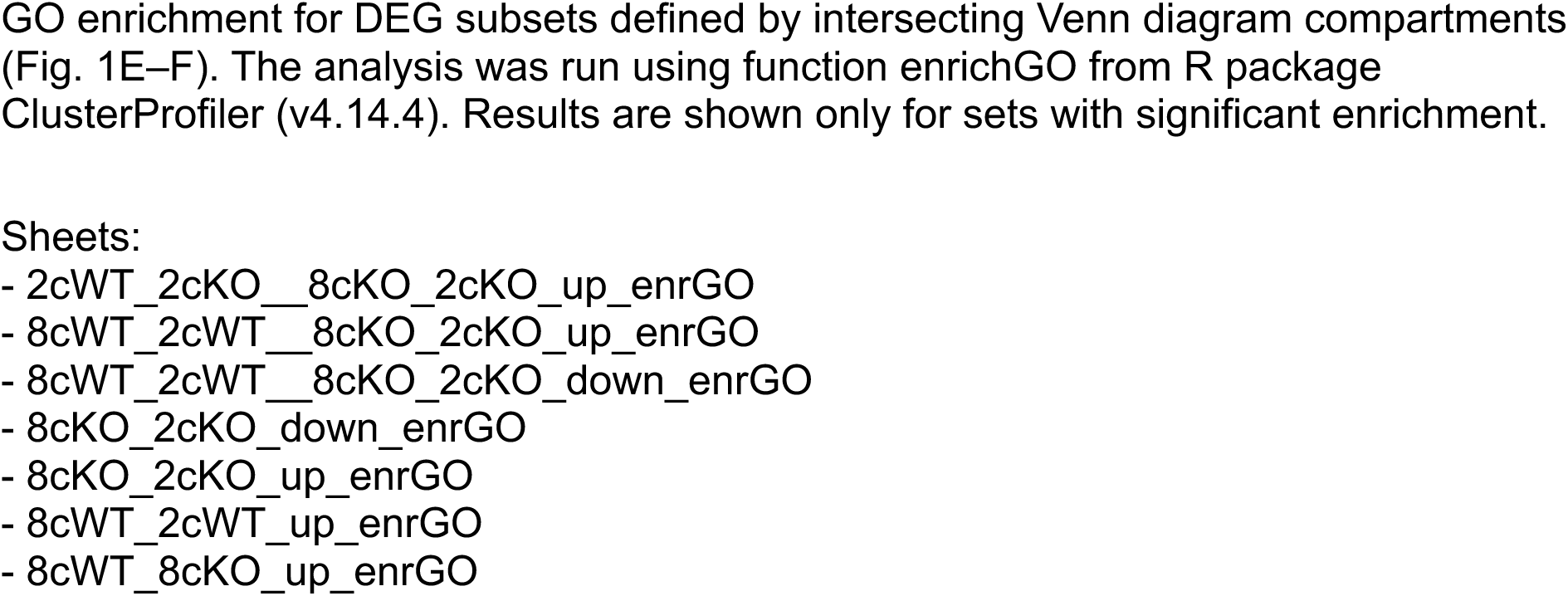
Overrepresentation Analysis (ORA) against Gene Ontology (GO) Analysis by DEG Venn Compartment.

**Table S5.**
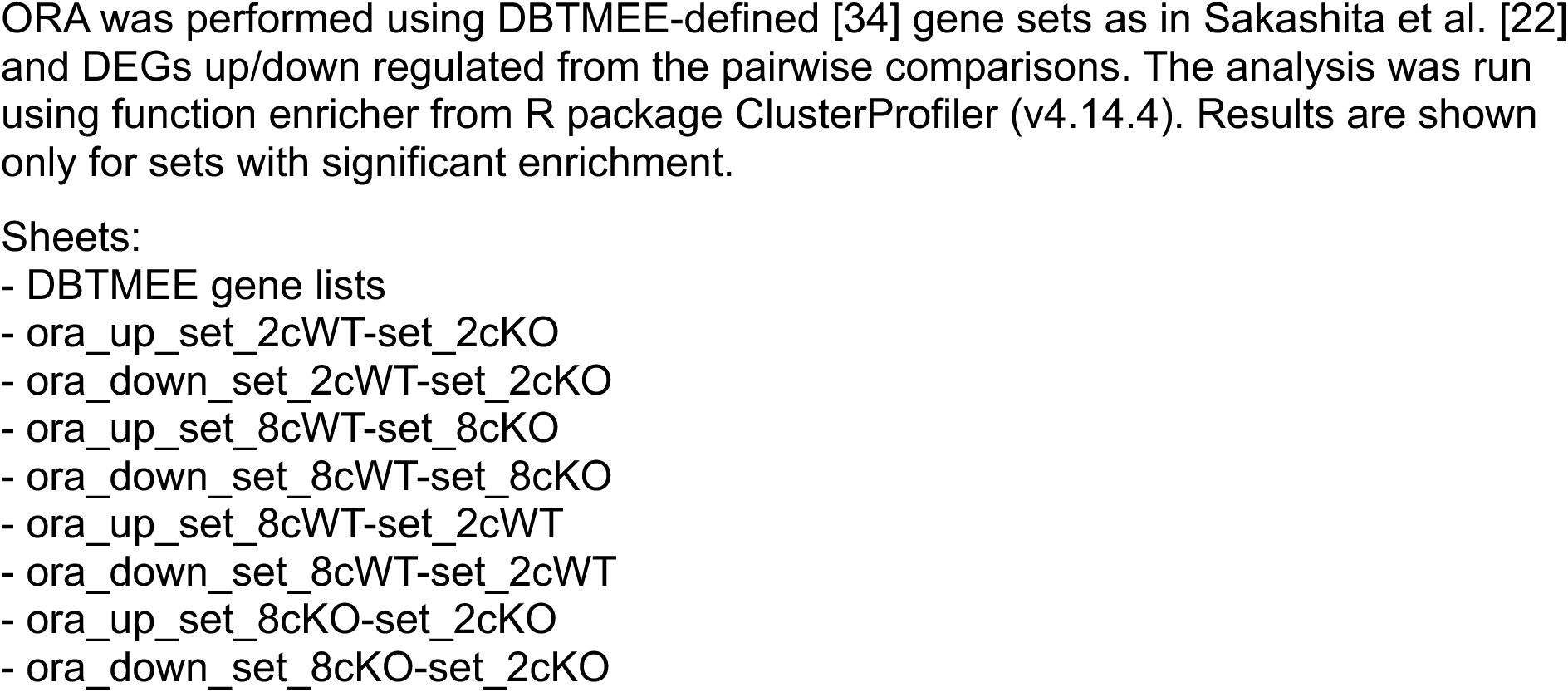
Overrepresentation Analysis (ORA) with DBTMEE Categories.

**Table S6.**
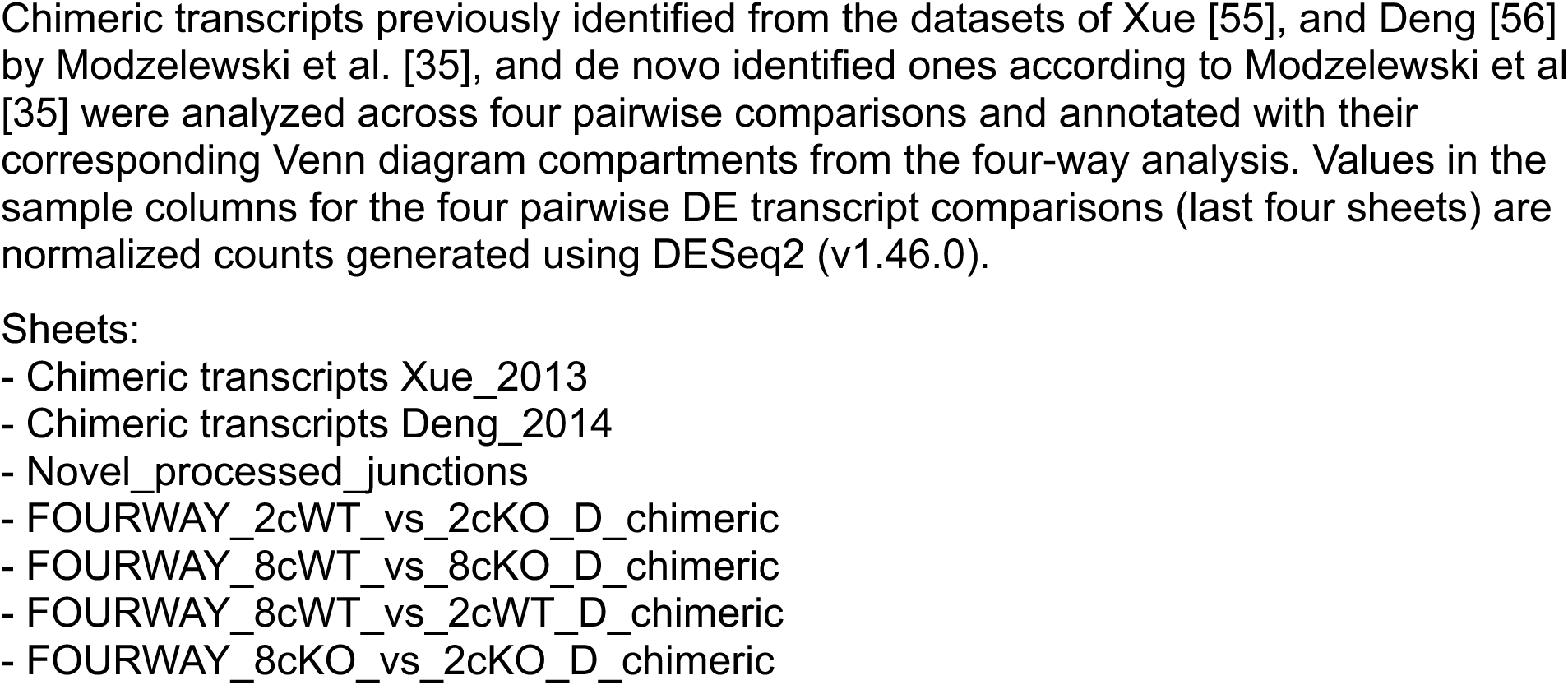
Chimeric Transcript Analysis.

**Table S7.**
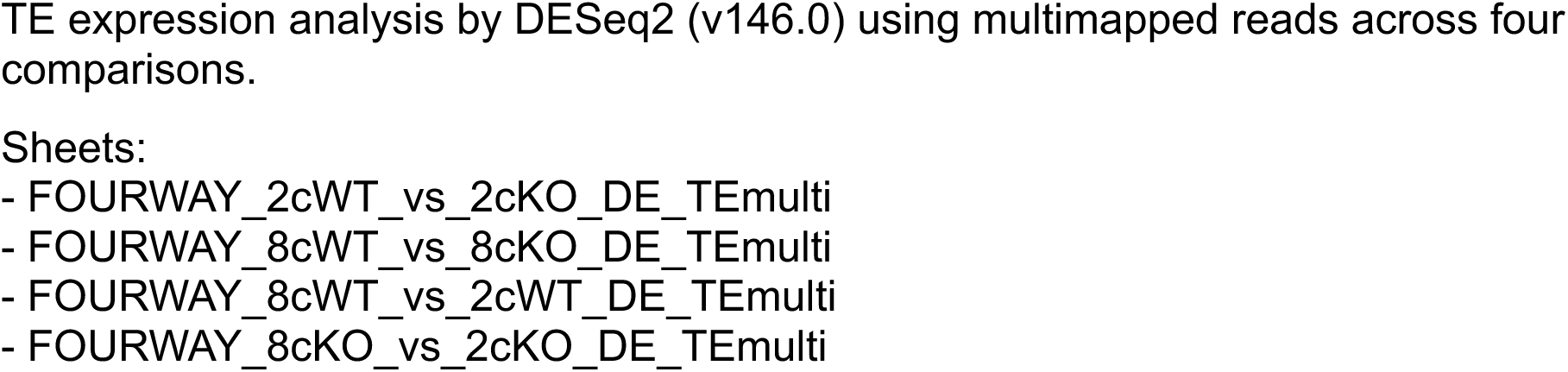
Differential TE Expression (DTE) – Multimapping Reads.

**Table S8.**
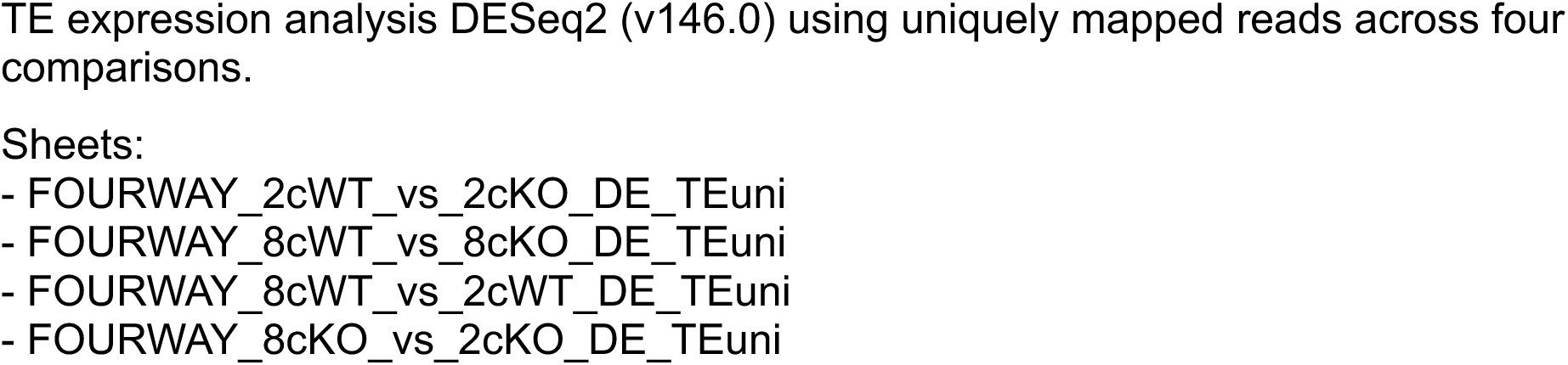
Differential TE Expression (DTE) – Unimapping Reads.

**Table S9.**
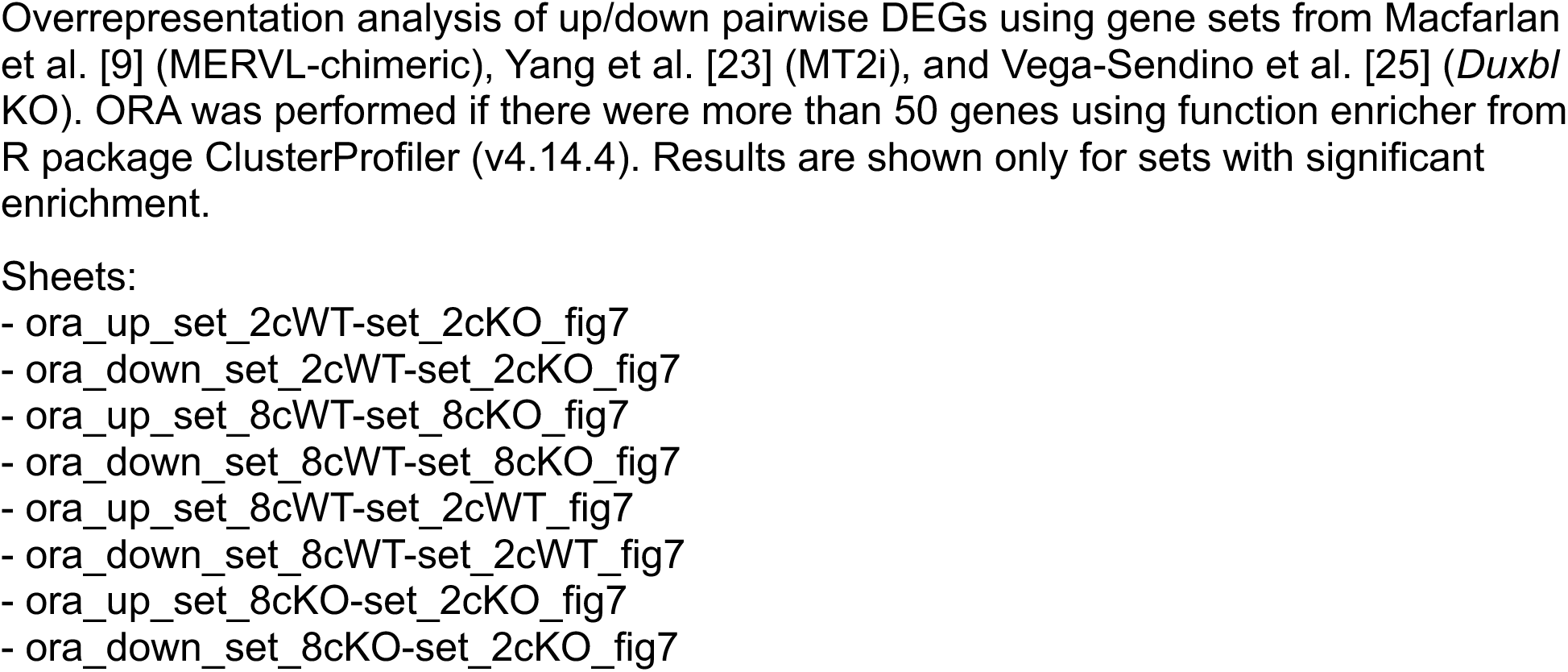
ORA with MERVL-Chimeric, MT2i, and *Duxbl* KO Gene Sets.

## Notes

### Competing Interest Statement

The authors have declared no competing interest.

